# B cell αv integrin regulates tissue specialization and clonal expansion of lung germinal center and memory B cells after viral infection

**DOI:** 10.1101/2024.04.03.587969

**Authors:** Andrea Montiel-Armendariz, Kelsey Roe, Jonathan Lagos-Orellana, Laura Veronica Martinez-Castro, Matthew J Dufort, Oliver Harrison, Adam Lacy-Hulbert, Mridu Acharya

**Author notes:** Corresponding author: Mridu Acharya, Seattle Children’s Research Institute, Seattle, WA, USA. AMA and KR contributed equally to the manuscript.

## Abstract

Lung-resident B cells are increasingly recognized as key contributors to protective immunity against respiratory viruses, yet the mechanisms that govern their generation and specialization remain poorly understood. Here, we identify B cell-intrinsic αv integrin as a critical negative regulator of germinal center (GC) dynamics and memory B cell formation in the lung following influenza A virus (IAV) infection. Using B cell-specific αv knockout mice, we show that loss of B cell αv integrin leads to persistent GC activity within the inducible bronchus-associated lymphoid tissue and expansion of lung-resident memory B cells, including IgA^+^ and cross-reactive B cells capable of recognizing heterologous influenza variants. Single-cell transcriptomic and B cell receptor (BCR) sequence analyses reveal that αv restricts clonal expansion and antigenic diversification of GC and memory B cells in the lung, but not in draining lymph nodes, indicating a spatially restricted mechanism of mucosal B cell regulation. These findings position αv integrin as a key checkpoint that constrains local mucosal B cell evolution and suggest new strategies to improve mucosal vaccine efficacy by enhancing GC activity directly in the lung.

## Introduction

Protective immunity against rapidly mutating pathogens such as respiratory viruses, HIV, and malaria depends on the ability of B cells to generate antibodies with high affinity and broad specificities as well as the long-term protection provided by memory B cells and long-lived plasma cells. While these features of B cell responses have been extensively studied in the context of systemic immune responses, recent studies are revealing the importance of localized B cell responses in mucosal tissues, particularly the lung, in combating respiratory pathogens.

Following influenza infection, early extrafollicular B cell responses are known to contribute to viral clearance, [1], while lung-resident memory B cells are known to provide critical protection against secondary infection by producing broadly reactive and often IgA-class–switched antibodies at mucosal sites[2–4]. IgA antibody, in particular, offers advantages due to its polymeric structure and ability to neutralize pathogens at epithelial barriers. However, despite growing evidence of their protective function, the mechanisms that control the generation and maintenance of these lung-resident B cells are still poorly understood. Moreover, these lung-resident B cells are not effectively induced by systemic vaccination strategies. Therefore, understanding the regulatory mechanisms behind lung-resident B cell responses is essential for developing vaccines that provide durable mucosal immunity against respiratory pathogens.

We previously identified a novel regulatory role for the αv integrin family in systemic B cell responses. αv integrins play diverse roles in multiple immune cell populations, such as dendritic cells, macrophages, T cells, and B cells[5–12]. We described a new function for αv integrin in regulating B cell responses to Toll-like receptor (TLR) ligands and TLR ligand-containing antigens, via regulation of intracellular trafficking events[13]. Mechanistically, αv integrin engages autophagy-related proteins to regulate the intracellular trafficking of TLRs and their ligands, thereby modulating the magnitude and duration of TLR signaling in B cells. Therefore, B cells lacking αv integrin show increased and persistent TLR signaling resulting in increased B cell responses to TLR-ligands[13, 14]. While αv-mediated regulation of TLR signaling serves to limit autoreactive responses in autoimmune settings[15], this same pathway also constrains protective immunity. Mice lacking B cell αv integrins develop heightened germinal center (GC) responses and show increased somatic hypermutation (SHM) of antibodies following systemic immunization with TLR ligand-containing antigens[16]. B cell specific αv knockout mice also showed better protection after infection with influenza virus [16]. However, it remained unclear whether this integrin-TLR axis influences B cell responses in mucosal tissues, where local inflammatory cues and persistent antigen exposure may uniquely shape immune regulation. Based on the known role of TLR signaling in mucosal B cell activation and the heightened presence of TLR ligands in inflamed lungs during infection, we hypothesized that B cell-intrinsic αv integrins may restrain local B cell responses in the respiratory tract.

To test this, we used a mouse model of intranasal (i.n.) IAV infection in combination with B cell-specific αv deletion, single-cell RNA sequencing, antigen-specific B cell tracking, and clonal lineage analysis. This approach allowed us to uncover a previously unrecognized, tissue-restricted role for αv integrin in regulating lung GC persistence, clonal expansion of B cells in the lungs, and the emergence of IgA^+^ and cross-reactive memory B cells. These findings reveal αv integrin as a key checkpoint that constrains mucosal B cell evolution and suggest new avenues for enhancing lung-resident humoral immunity through targeted modulation of local GC activity.

## Results

### B cell αv integrin regulates generation of cross-reactive lung-resident GC and memory B cells following influenza infection

To investigate whether B cell-intrinsic αv integrin regulates tissue-specific B cell responses during respiratory viral infection, we intranasally infected B cell specific αv conditional knockout mice (referred to as cKO mice) and matched controls with a sub-lethal dose of H1N1 PR8 IAV (**Fig 1A**). Fourteen days post-infection, we used ELISPOT assay to identify overall changes in antigen specific B cells from lung and draining mediastinal lymph nodes (medLNs). Compared to controls lungs from cKO mice showed increased numbers of PR8-HA–specific IgG-secreting B cells after infection (**Fig 1B**). Moreover, Cal09-HA-specific IgG-secreting cells were significantly increased in cKO lungs compared with controls (**Fig 1B**). In contrast, numbers of HA-specific antibody-secreting cells in the medLN were similar between groups (**Fig 1C**), suggesting that αv deficiency specifically enhances local lung B cell responses after infection.

**Figure 1.**
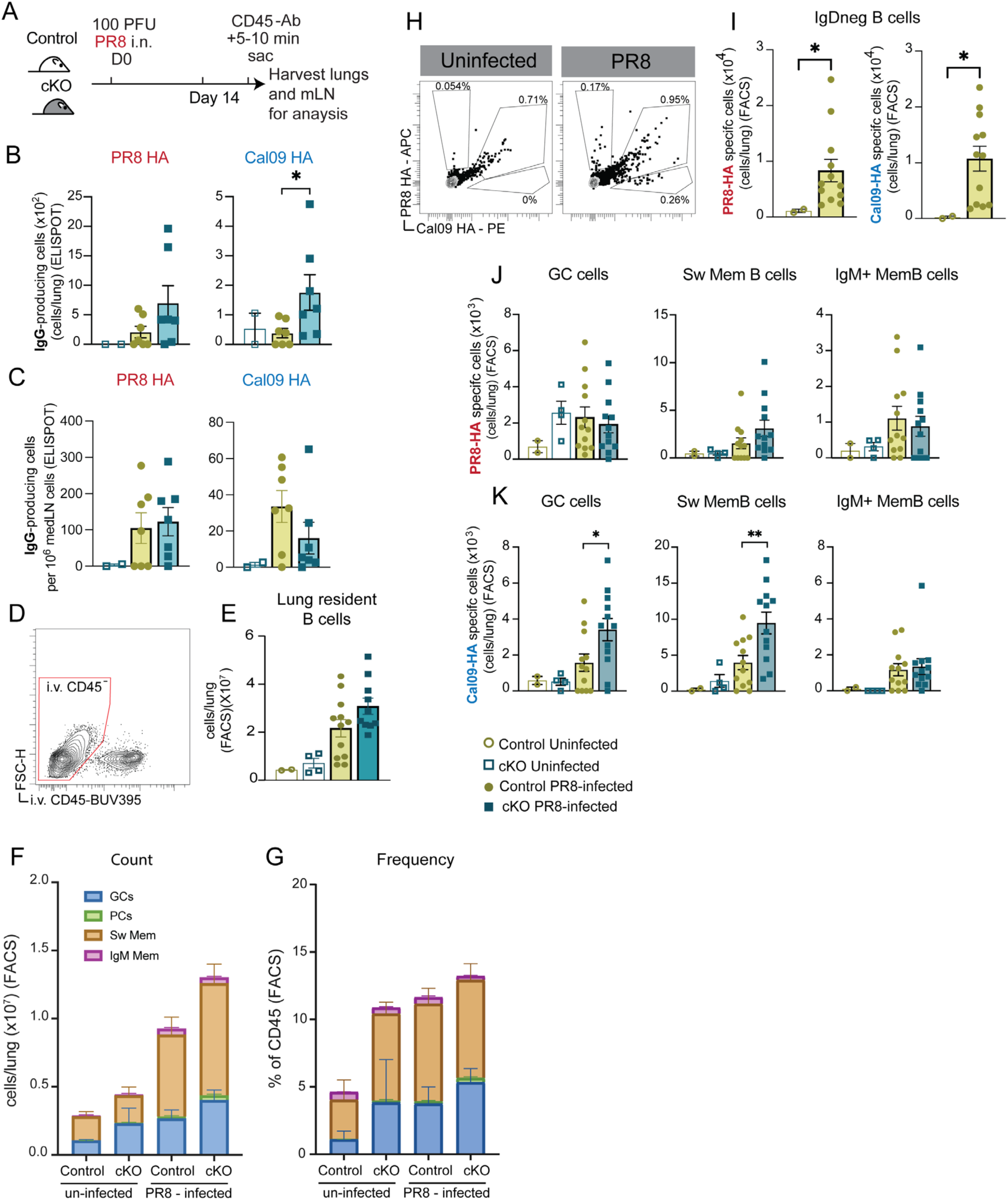
Generation of cross-reactive B cell responses in the lungs is regulated by B cell αv integrin. αv^+/+^ CD19^Cre+^ (control) and αv^fl/fl^ CD19^Cre+^ (cKO) mice were infected intranasally (i.n.) with 100 plaque-forming units (PFU) of live PR8 IAV. 14 days later all mice received a retro-orbital (r.o.) injection with 1µg anti-CD45 antibody (Ab) (α-CD45-BUV395) five minutes prior to euthanasia. **(A)** Schematic for experimental procedure. **(B - C)** Quantification of IgG-secreting cells in the lungs **(B)** or medLN **(C)**, that recognize either PR8-HA (left) or Cal09-HA (right) as identified by ELISPOT. **(D)** Representative flow cytometry plot of the i.v. CD45 staining in the lungs. **(E)** Quantification of live B cells gated as lung resident (i.v. CD45-unlabeled) B cells (CD19^+^ CD138^+/-^). **(F - G)** Count **(F)** and frequency (% of i.v. CD45-unlabeled cells, **G**) of B cell subpopulations. Lung resident B cells gated into sub populations based on the following surface markers: Plasma cells (PCs) were gated as IgD^-^ IRF4^+^ CD138^+^ cells, germinal center cells (GCs) were gated as IgD^-^ IRF4^-^ CD138^-^ Fas^+^ PNA^+^ and Sw memory B cells were gated as IgD^-^ IRF4^-^ CD138^-^ Fas^-^ PNA^-^ IgM^-^ IgA^-/+^ (see **Supplemental** Figure 1 for gating strategy).Cells per lung were calculated by multiplying the frequency of each population (% of live cells), as determined by flow cytometry, by the number of total live cells recovered from the tissue harvest. **(H)** Representative flow plot of PR8-HA – APC and Cal09-HA – PE tetramer staining in an uninfected (left) and infected mouse (right) on lung resident IgD^-^ B cells. **(I)** Quantification of lung resident IgD^-^ B cells specific for PR8-HA (left) and Cal09-HA (right). **(J - K)** Quantification of lung resident PR8-HA specific **(J)** or Cal09-HA specific **(K)** GCs, Sw Memory B cells and IgM+ Memory B cells. Each dot represents an individual mouse (n= 2-4 mice for uninfected group and n=7-12 mice for infected groups). Data are means ± SEM of either the combination of two independent experiments **(B – I)** or a representative experiment **(J - K)**. \**p*<0.05, \*\**p*<0.01 by Mann-Whitney U-test between the two PR8-infected groups.

To determine which B cell subsets accounted for these tissue-specific changes, we used flow cytometry with intravenous (i.v.) labeling to distinguish circulating B cells from lung-resident B cells and gated on different B cell subpopulations (**Supp Fig 1**; **Fig 1D**). Total numbers of i.v.^-^ lung-resident B cells increased equivalently post-infection in both genotypes (**Fig 1E**), and this increase was largely driven by memory B cells accumulation (**Fig 1F-G**). We also identified i.v.^-^ lung-resident GC-like B cells by surface expression of FAS^+^ PNA^+^, which were present at higher baseline levels in cKO lungs, consistent with our prior observations of increased spontaneous GCs in Peyer’s patches [16]. These lung-resident GC-like B cells were expanded upon infection in both genotypes with the largest increase in the cKO mice, suggesting a role for αv in constraining GC responses in the lungs (**Fig 1F-G**).

We next evaluated the antigen specificity and cross-reactivity of the lung-resident B cell subpopulation using hemagglutinin (HA) tetramers from PR8 and Cal09 IAV strains [17]. After verifying these tetramers could identify lung-resident PR8-HA^+^ or Cal09-HA^+^ B cells induced by PR8 infection (**Fig 1H-I**), we compared HA-specific lung-resident B cells in the PR8 infected control and cKO mice. The number of infection-induced PR8-HA-specific lung-resident GC B cells was not significantly altered between the genotypes; although, class-switched memory B cells were slightly increased in cKO lungs (**Fig 1J**). In contrast, infection induced lung-resident Cal09-HA-binding B cells of GC and memory B cell phenotype were significantly elevated in cKO lungs compared to controls (**Fig 1K**), indicating enhanced cross-reactive responses in the absence of αv. Furthermore, lung-resident IgM^+^ memory B cells specific for either PR8-HA or Cal09-HA were not increased in cKO lungs, suggesting that αv deficiency preferentially enhances class-switched antigen-experienced subsets (**Fig 1 J-K**). These differences were not observed in the medLN, where HA-specific GC and memory B cell frequencies were similar between groups (**Supp Fig. 2**), reinforcing the tissue-restricted effect of B cell αv loss in promoting B cell responses after intranasal infection.

**Figure 2.**
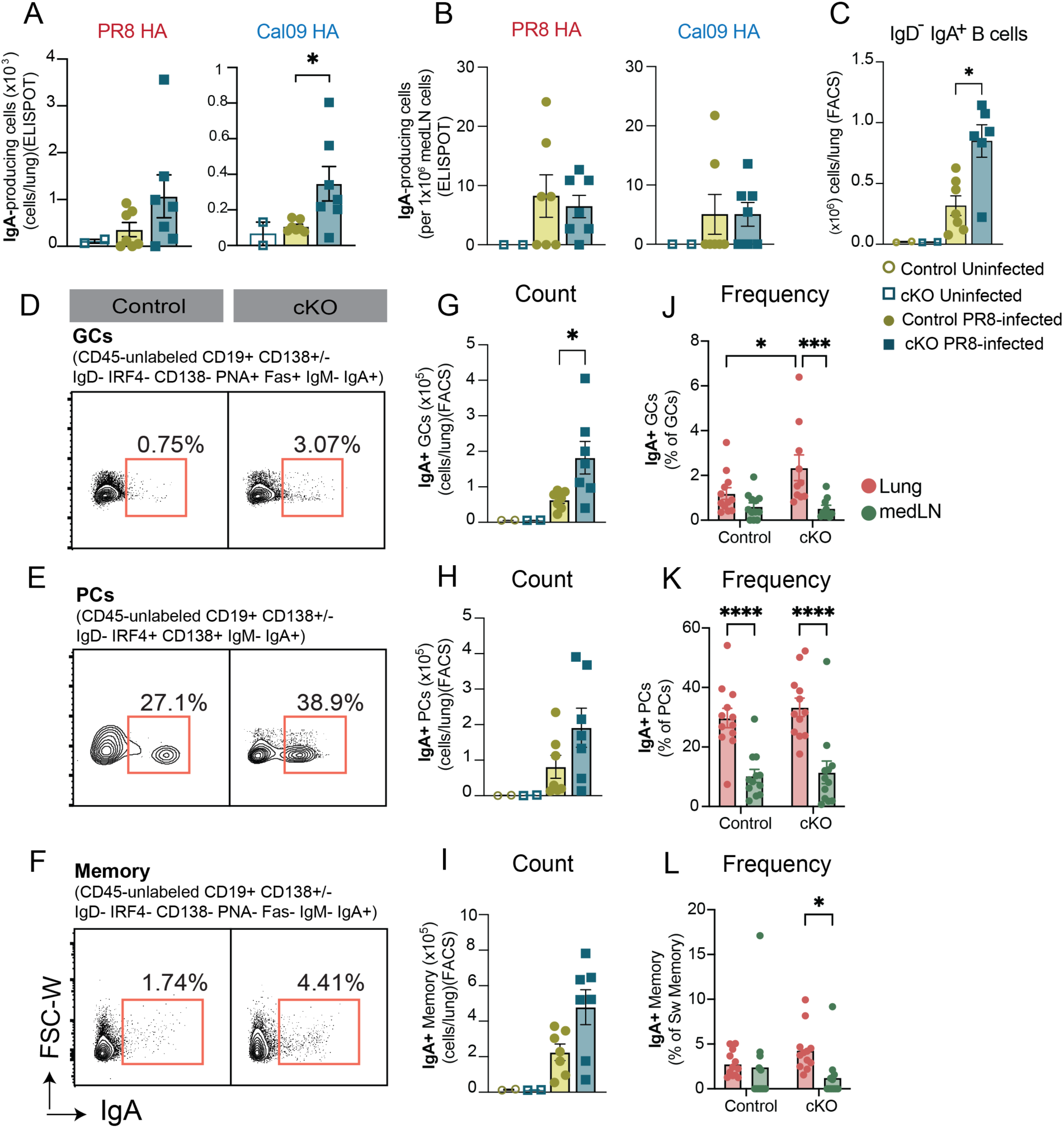
Absence of B cell αv integrin promotes IgA B cell response in the lungs after influenza infection. αv^+/+^ CD19^Cre+^ (control) and αv^fl/fl^ CD19^Cre+^ (cKO) mice were infected i.n. with 100 plaque-forming units (PFU) of live H1N1 PR8 IAV. After 14 days of infection all mice received r.o. injection with 1µg α-CD45-BUV395 five minutes prior to euthanasia. (**A-B**) IgA-secreting cells in the lungs **(A)** or medLN **(B)** that recognize PR8-HA (left) or Cal09-HA (right) as detected by ELISPOT. **(C)** Quantification of lung resident IgD^-^ IgA^+^ B cells by flow cytometry. **(D-F)** Representative flow plots of the expression of IgA on GC B cells **(D)**, PCs **(E)** and mem B cells **(F)** (See Supplemental Figure 1 for gating strategies). **(G-I)** Quantification from flow analysis of IgA+ GCs **(G)**, PCs **(H)**, and IgA-memory B cells **(I)**. **(J-L)** Quantification of the frequency of IgA+ GCs **(J)**, PCs **(K)** and memory **(L)** as a frequency of their respective parent population in lungs and medLN. Each dot represents an individual mouse (n= 2mice for uninfected group and n=6-7 mice for infected groups). Data are means ± SEM from one representative experiment from 3 independent experiments. \**p*<0.05, ***p<0.001, ****p<0.0001 by Mann-Whitney U-test between the two PR8-infected groups **(A-I)** or 2way ANOVA comparing tissue and group **(J-L)**.

These findings demonstrate that B cell-intrinsic αv integrin specifically restrains the generation of cross-reactive, lung-resident, GC and memory B cells following influenza infection.

### Loss of B cell αv integrin increases lung-resident IgA+ B cells following influenza virus infection

Secretory IgA plays a key role in mucosal defense against respiratory viruses, but the mechanisms governing generation of IgA-class-switched B cells in the lung remain poorly understood. Based on the enhanced lung B cell responses seen in cKO mice, we examined whether B cell-intrinsic αv integrin influences the generation of IgA^+^ B cells following influenza infection.

ELISPOT analysis of HA-specific cells showed that at day 14 post infection, both cKO and control mice generated HA-reactive IgA-producing cells in the lungs (**Fig 2A**). Anti-PR8-HA IgA-producing cells were present at higher numbers in some cKO mice compared with controls, but this did not reach statistical significance. However, Cal09-HA-binding cells-indicative of cross-reactive responses-were more consistently increased in cKO mice compared with controls, similar to our findings for IgG-producing cells (**Fig 2A**). In contrast to the lung, the number of HA specific IgA-secreting cells was similar between groups in the medLN (**Fig 2B**) suggesting B cell αv deficiency specifically enhances lung-localized IgA responses.

To characterize these IgA^+^ B cells in the lungs, we performed flow cytometric phenotyping of lung-resident B cells. IgD⁻ IgA⁺ B cells were rare prior to infection in both genotypes, but were robustly induced in the lungs following infection, with cKO mice displaying a ∼2-fold increase in total lung-resident IgA^+^ B cells compared to controls (**Fig 2C**). This increase extended across the GC, plasma cell (PC), and class-switched memory B cell compartments (**Fig 2D–I**), indicating that αv deficiency promotes IgA class switching related to multiple B cell subsets. In contrast, no significant genotype-dependent differences in IgD^-^ IgA^+^ B cells were observed in the medLN (**Supp Fig 3A**).

**Figure 3.**
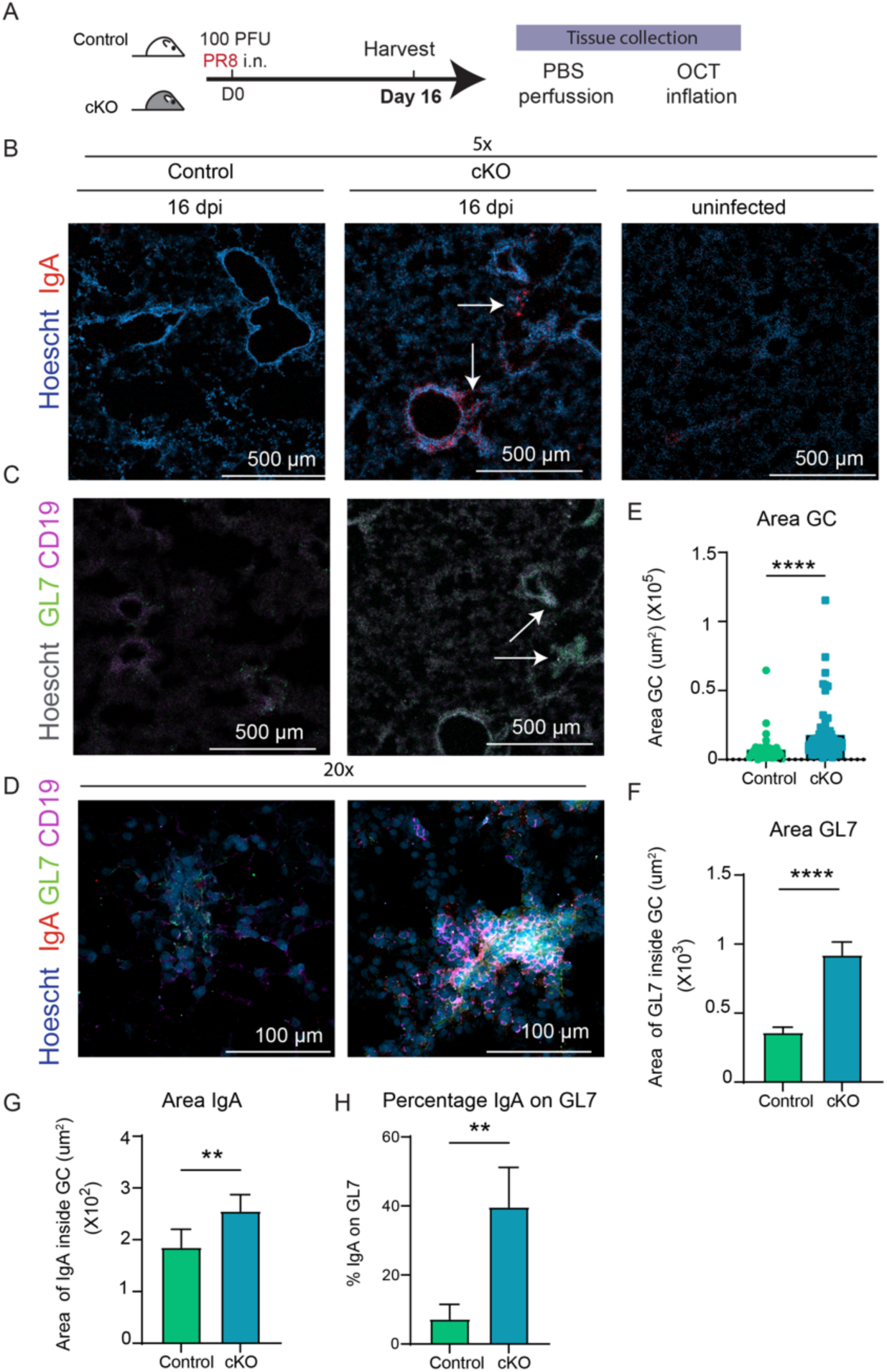
Loss of B cell αv integrin enhances the induction of iBALT in lungs after viral infection. αv^+/+^ CD19^Cre+^ (control) and αv^fl/fl^ CD19^Cre+^ (cKO) mice were infected i.n. with 100 plaque-forming units (PFU) of live PR8 IAV. Lungs were harvested after 16 days post infection (dpi). (**A**) Schematic of mice infection and tissue collection for lung tissue staining. (**B-C**) Representative confocal images of lungs from uninfected control mouse and infected control or cKO mice. Images representing one single plane at 5x magnification. Scale bar 500 µm. (**B**) Lung staining for Hoechst (nucleus, Blue) or IgA (red). Airways surrounded by IgA staining are shown by arrows. (**C**) Lung staining for Hoechst (nucleus, gray), GL7 (green), and CD19 (magenta). Arrows represent GL7 positive and CD19 positive structures corresponding to iBALTs. (**D**) Representative confocal images of lungs from control or cKO mice 16 dpi. Images representing one single plane at 20x magnification labeled with Hoechst (nucleus, Blue), IgA (red), GL7 (green), and CD19 (magenta) showing iBALTs. Scale bar 100 µm. (**E**) Scatter plot with a bar graph showing the average size (µm^2^) of GL7+CD19+ iBALT structures by manually quantifying the area of these structures. Every dot is the size of 1 iBALT structure (35-65 structures) from 22-28 different fields of sections viewed at 10x from 3 different mice per genotype. (**F and G**) Bar graph showing mean and SEM of the area (µm^2^) of (**F**) GL7 and (**G**) IgA positive staining inside the structures (N= 35-65) as defined in **E**. (**H**) Bar graph showing mean and SEM of the percentage of IgA+GL7+ staining within iBALT structures. Sections (N=7-9) viewed at 20x from 3 different mice per group. **p<0.01, ****p<0,0001 by Mann-Whitney U-test.

We next compared the distribution of IgA^+^ B cells between the medLN and lung tissue. While both sites showed infection-induced IgA^+^ B cells, there was a higher proportion of IgA^+^ cells within the lung-resident B cell compartments in both genotypes (**Fig J-L**). Moreover, the cKO mice showed a significant increase in the frequency of IgA^+^ cells within the lung-resident GC B cell compartment. Meanwhile the frequency of IgA^+^ B cells in the medLN remained low in both genotypes across all B cell compartments (**Fig J-L**). Consistent with this, comparison of IgA vs IgG secreting HA-specific cells in the lungs and medLN also showed that IgA-secreting cells were found predominantly in the lungs compared to the medLN and the increase observed in cKO mice was predominantly localized to the lungs (**Sup** Figure 3B-C).

Taken together, these data demonstrate that IgA^+^ B cells preferentially accumulate in the lung following influenza infection, and that B cell-intrinsic αv integrin limits the generation of these cells, particularly those with a GC phenotype.

### Loss of B cell αv integrin promotes infection induced GC reactions locally in the lungs

To investigate whether the loss of B cell αv promotes lung-resident IgA^+^ B cells via local GC activity, we used confocal microscopy to examine lung sections from infected mice at day 16 post-infection (**Fig 3A**). We confirmed that more IgA^+^ cells were present in cKO lungs compared to controls, and these cells were distributed around large airways and within clusters in the lung parenchyma (**Fig 3B**). We therefore asked whether these clusters of IgA^+^ cells were related to GC structures in the lungs.

Ectopic lung GC reactions have previously been shown to develop in inducible bronchus-associated lymphoid tissues (iBALT) after influenza infection [18, 19]. We therefore used CD19 staining to identify B cells and GL7 for GC B cells (**Fig 3C**; **Supp** Fig 4) and manually defined the regions where GL7^+^ B cells clustered together as the iBALT-GCs induced by infection. These iBALT regions with GC B cells were close to airways and found in the lungs of both control and cKO mice. Although the iBALT regions were not as abundant at this time point, as described by others at later time points[20], the cKO lungs exhibited larger and more organized clusters of GC B cells, whereas control lungs showed smaller, less organized GC clusters. (**Fig 3C and D**). Quantification of either the manually gated iBALT-GC area (Area GC) or levels of GL7 staining within the iBALT regions both confirmed that the clusters of GC cells were larger in the lungs from cKO mice, compared to the lungs of the control mice (**Fig 3E-F**).

**Figure 4.**
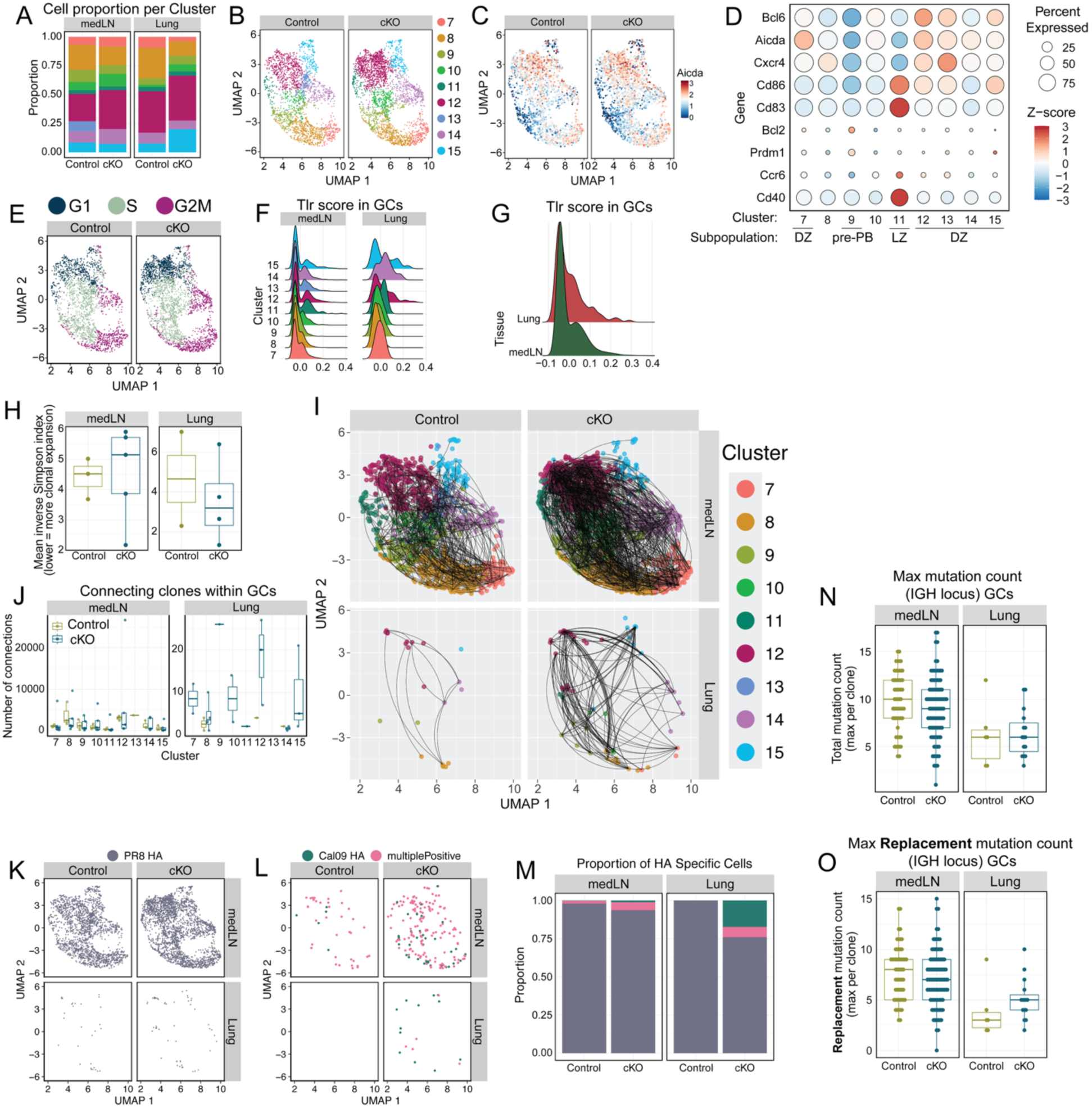
Loss of B cell av leads to expansion of GC sub-clusters in the lungs and promotes cross-reactivity. Single cell RNA sequencing analysis of sorted lung resident and medLN antigen specific (PR8-HA+) and cross reactive (Cal09-HA+) B cells from αv^+/+^ CD19^Cre+^ (control) and αv^fl/fl^ CD19^Cre+^ (cKO) mice 20 days post i.n. infection with 100 plaque-forming units (PFU) of live PR8 IAV. (**A - B**) GC subclusters as identified by Louvain method. (**A**) Stacked bar graph of the proportion of cells in the GC subclusters in lung and medLN of the control and cKO mice. (**B**) UMAP of the GC subclusters in control and cKO mice. (**C**) Expression of *Aicda* in the control and cKO GCs. (**D**) Dot plot showing expression of key genes in GC subclusters; color represents the expression level (z-score), circle size represents proportion of cells expressing that marker. (**E**) UMAP of the GC cells colored by cell cycle phase as identified with Seurat (left). **(F - G)** Ridge plot of the *Tlr* score expression comparing by cluster (**F**) and by tissue (**G**) in the GC cells. Score obtained from the Seurat package by calculating the average expression of a gene set and subtracting the aggregated expression of control gene sets. Gene set for *Tlr* score: *Tlr-1, 2, 3, 4, 6, 7,* and *9*. **(H)** Inverse Simpson index of BCR clonal diversity; values are mean of repeated down sampling to a common number of GC cells per mouse in each tissue. **(I)** UMAP of the GC subclusters showing BCR clonal relationships. Lines connect cells from the same BCR clone within each tissue; to illustrate differences in clonal expansion, only 1/100 connections are shown. **(J)** Quantification of the number of connecting lines of a clone to its sister clones within the GCs. Dots represent outliers. **(K - L)** UMAP of the PR8-HA (**K**) and cross-reactive (**L**; Cal09-HA+ or multiple positive: binds to both PR8-HA and Cal09-HA tetramers) GCs. **(M)** Quantification of the proportion of cells by specificity in the medLN and lungs of the control and cKO mice. **(N - O)** Quantification of the maximum number of total (**N**) and amino acid-replacing mutations (**O**) by clone in total GCs in the medLN and lung of the control and cKO mice. Each dot represents one clone.

Moreover, IgA+ cells were more frequently localized to iBALT-GCs in cKO lungs than in controls (**Fig 3G**), and IgA/GL7 co-localization was significantly higher in cKO sections (**Fig 3H**). These imaging results were consistent with increased numbers of IgA^+^ GC B cells observed by flow cytometry (**Fig 2D and G**). These findings show that loss of B cell αv integrin promotes the formation of ectopic GC structures directly in the lungs and enhances the generation of GC derived IgA^+^ B cells following influenza infection.

### Loss of B cell αv leads to expansion of GC sub-clusters in the lungs and promotes cross-reactivity

The increase in the ectopic GC structures in the lungs of cKO mice raised questions about whether these GC cells were similar to canonical GC cells, such as those in the medLN. To address this question, we turned to single cell RNA sequencing (scRNAseq) of Ag-reactive B cells induced after infection. PR8-HA^+^ (Ag-specific) and Cal09-HA^+^ (cross-reactive) tissue-resident B cells were sorted from lungs and medLNs of cKO or control mice, 20 days after infection with H1N1 PR8. In these experiments, we made use of conformationally intact HA tetramers [21, 22] rather than the linear HA probes used in Fig 1, to more sensitively identify HA-specific B cells for analysis. We verified that these tetramers could be used to identify antigen-reactive cells by flow cytometry (**Sup Fig 6A**).

For each B cell we simultaneously obtained HA specificity (via oligo-tagged PR8-HA and Cal09-HA probes), BCR sequence from heavy and light chains, Ab isotype and transcriptome with the 10x Chromium technology (**Supp Fig 5A**). Clustering by Louvain community detection based on gene expression of all B cells identified 16 different clusters, which were assigned to one of four major cell types (naïve, GC, memory, and plasma cells) across both tissues based on the expression of *Ighd, Aicda, Cd80 and Prdm1* (**Supp Fig 5B**). Deconvolution of oligo hash-tagged antibodies showed that some mice lacked antigen-specific GC B cells in the medLN, which indicated unsuccessful influenza infection. Consequently, cells from these uninfected mice were removed from downstream analysis. The remaining 8 mice showed even representation of high-quality antigen-specific cells. We first focused on the GC clusters (clusters 7 – 15) and observed that lung-derived GC cells shared transcriptional similarities with those from the medLN. Despite being fewer in number, lung-derived GCs clustered together with comparable GC subtypes found in the medLN (**Supp Fig 5B-C**). Notably, all GC subclusters were represented in both lung and medLN, with the exception of cluster 13, which was only present in the medLN (Fig 4 **A-B**).

**Figure 5.**
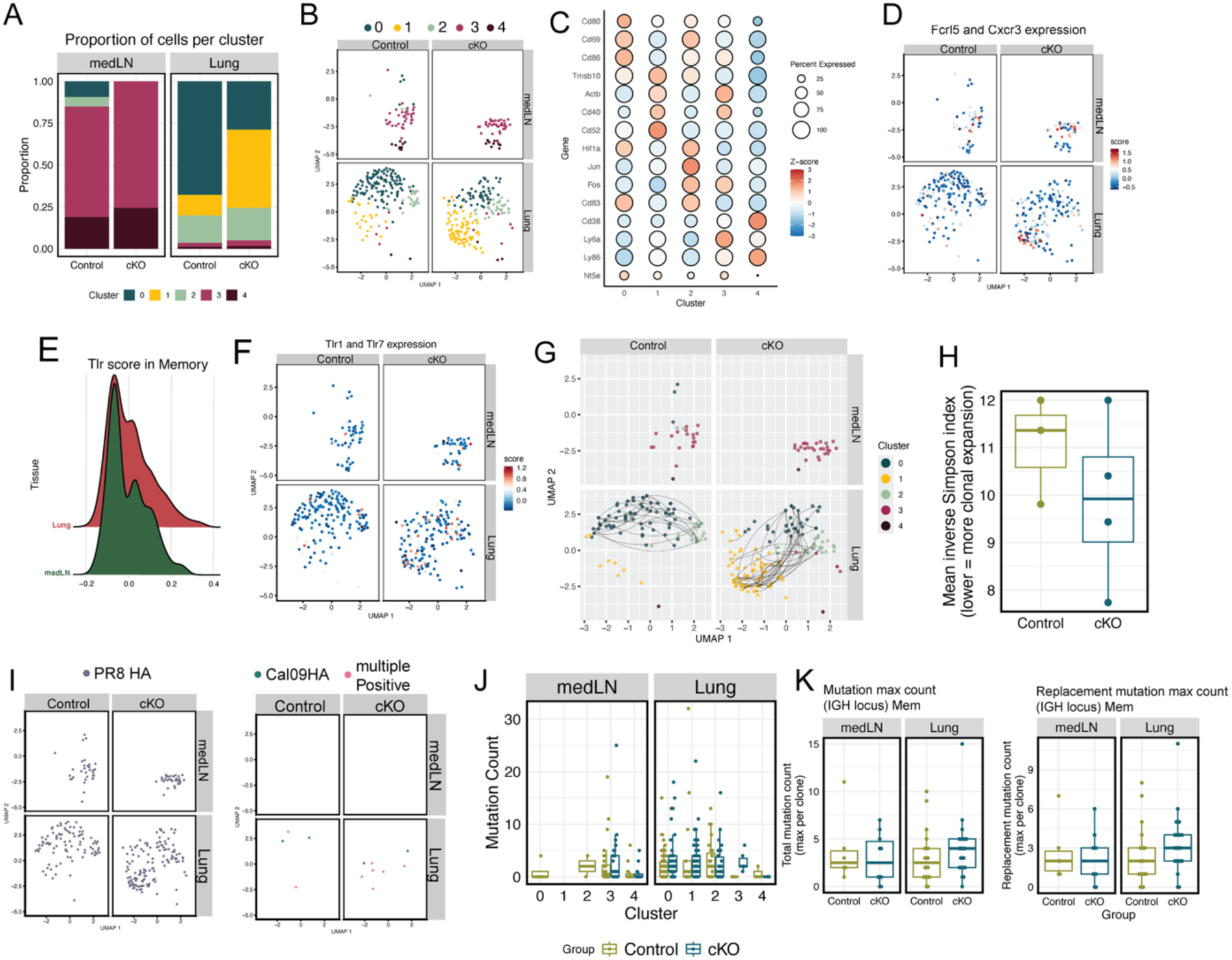
Loss of B cell av induces unique memory B cell subsets in the lungs. Single cell RNA sequencing analysis of sorted lung resident and medLN antigen specific (PR8-HA+) and cross reactive (Cal09-HA+) B cells from αv^+/+^ CD19^Cre+^ (control) and αv^fl/fl^ CD19^Cre+^ (cKO) mice 20 days post i.n. infection with 100 plaque-forming units (PFU) of live PR8 IAV. **(A-B)** Sub clustering of the Cd80+ memory B cell cluster, cluster 6. **(A)** Quantification of the proportion of cells and **(B)** UMAP representation of the distribution per cluster in the medLN and lungs of the control and cKO groups. **(C)** Expression of marker genes within the different memory subclusters; color represents the expression level (z-score) of genes and circle size the proportion of cells expressing that marker. **(D)** Expression of *Fcrl5* and *Cxcr3* on memory cells (calculated as in **Fig 4G**) in medLN and lungs of control and cKO groups. **(E)** Distribution of TLR score (calculated as in **Fig 4G**). **(F)** Expression score of *Tlr1* and *Tlr7* in the memory B cell subset. **(G)** Clonal sharing by BCR sequences, comparing clonality between memory cell subclusters in the medLN and lungs of the control and cKO group. Each dot represents a cell; each color represents a subcluster and each line connects a shared clone. **(H)** Clonal diversity of lung memory cells from control and cKO mice, quantified by inverse Simpson index; values are means of 100 replicates downsampled to a common number of lung memory cells per mouse. **(I)** UMAP representation of the PR8-HA specific (left) and cross reactive (Cal09-HA+ and multiplePositive: PR8-HA+/Cal09-HA+) in the memory subclusters. **(J-K)** Quantification of the maximum number of total mutations by clone in memory subclusters **(J)** and in total memory cells (**K**, left) and maximum number of replacement mutations in memory cells (**K**, right) in the medLN and lung of the control and cKO mice. Each dot represents one clone.

**Figure 6.**
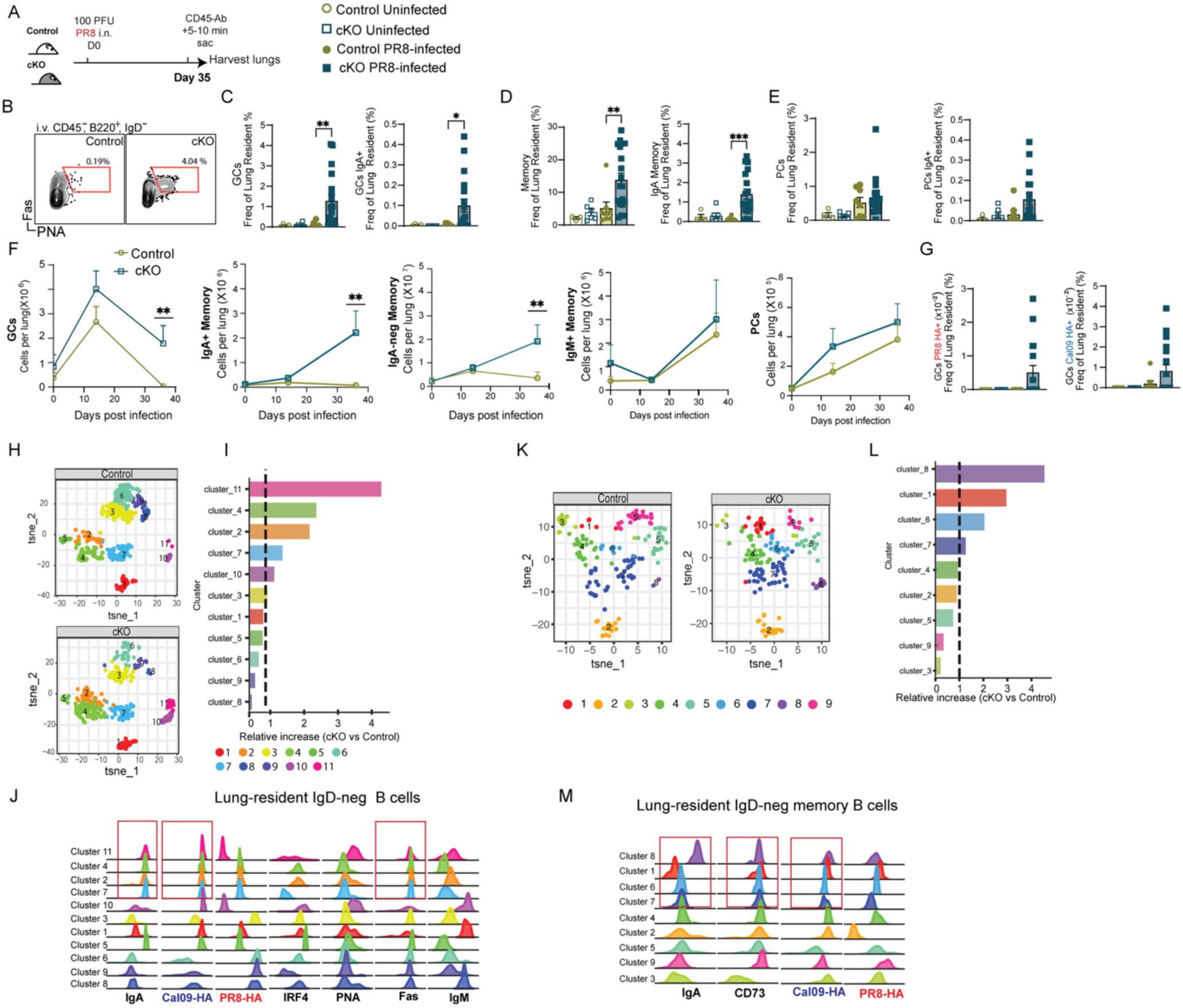
Absence of B cell αv integrin promotes sustained increase in GC and memory B cell responses in the lungs after infection. αv^+/+^ CD19^Cre+^ (control) and αv^fl/fl^ CD19^Cre+^ (cKO) mice were infected intranasally (i.n.) with 100 plaque-forming units (PFU) of live PR8 IAV. 35 days later all mice received a retro-orbital (r.o.) injection with 1µg anti-CD45 antibody (Ab) (α-CD45-APC) five minutes prior to euthanasia. **(A)** Schematic for experimental procedure. **(B)** Representative flow plot of the GC cells; gated as resident (CD45-unlabeled) B220^+^ IgD^-^ IRF4^-^ Fas^+^ PNA^+^ from a control (left) and a cKO (right) mouse. **(C-E)** Quantification of the frequency of total GCs and IgA+ GCs **(C)**, Sw Memory and IgA+ Memory **(D)** and PCs and IgA+ PCs **(E).** Each dot represents one mouse, reported as frequency of resident lung cells; data are means SEM± of one representative experiment from three independent repeats. *p<0.05, **p<0.001, ***p<0.0001 by Mann-Whitney U-test comparing the two PR8 infected groups. **(F)** Line graph of the number of resident lung GCs, IgA+ Memory, IgAneg Memory (Sw Memory), IgM+ Memory, and PCs. Cell number was obtained as in Fig 1. **(G)** Quantification of the frequency of lung resident GC specific for PR8-HA (left) or Cal09-HA (right) as identified by flow cytometry. **(H)** t-SNE plot of HA-specific lung-resident IgD-B cells (i.v CD45-B220+ IRF4+/- IgD-PR8-HA+/Cal09-HA+) from lungs of infected control (top) and cKO mice (bottom) with the PhenoGraph clusters (left). **(I)** Fold increase of the cell proportion per cluster in the cKO vs the control mice (right). **(J)** Histograms of the expression of the markers included in the analysis for each cluster, red boxes include the clusters with an increased frequency in the cKO mice compared to the control (bottom). (**K**) Mice were infected i.n. with 100 plaque-forming units (PFU) of live PR8 IAV and re-challenged at day 31 with 50 PFU live PR8. All mice received r.o. injection with 1µg α-CD45-APC five minutes prior to euthanasia. t-SNE plot of lung-resident IgD-memory B cells (iv CD45-B220+ IRF4+/- IgD-, Fas-PNA-) from infected control (left) and cKO mice (middle) with the PhenoGraph clusters. **(L)** Fold increase of the cell proportion per cluster in the cKO vs the control mice (right). **(M)** Histograms of the expression of the markers included in the analysis for each cluster, red boxes include the clusters with an increased frequency in the cKO mice compared to the control (bottom). Analysis is based on n= 6 mice per group in both sets of cluster analysis.

Sub-clustering of GC B cells (**Fig 4A-B**) identified expected subpopulations consistent with known GC biology, including subclusters showing higher *Aicda* expression (**Fig 4C**). We identified dark zone (DZ)-like subclusters which showed higher expression of *Cxcr4, Aicda* and were *Bcl6^+^ (clusters 7, 12-15)* as well as clusters showing increased expression of markers consistent with light zone (LZ) cells (*Cd83^+^*, *Cd86^+^*or *Cd40^+^*, cluster 11), and putative pre-plasmablast cluster (*Bcl6^+^, Prdm1^+^,* cluster 9) (**Fig 4D**). Further analysis of cell cycle gene expression within these GC subclusters also revealed dynamics consistent with genuine GC reactions (**Fig 4E**). Specifically, DZ subcluster 12 showed an increased proportion of cells in G1 phase and high *Aicda* expression, aligning with the expected characteristics of DZ cells that are undergoing SHM [23, 24]. Moreover, other DZ-like clusters showed distinct cell cycle phases, with subcluster 7, 14, and 15 undergoing G2/M phase and subcluster 13 in S phase (**Fig 4E and Supp Fig 5D**). Hence, we concluded that the lung GCs represented genuine GC cells, with transcriptional patterns consistent with DZ cells undergoing proliferation and SHM (**Fig 4E; Supp Fig 5D**).

Importantly, GC cells from cKO mice showed selective expansion of several DZ-like subclusters (clusters 7, 12, 14, 15) (**Fig 4A-B; Supp Fig 5D**). These expanded clusters included those enriched for cells in G1 phase (clusters 12,15), a stage associated with SHM [23], and proliferating cells in G2/M phase (clusters 7,14) (**Fig 4E and Supp Fig 5E**). These data suggested that αv deficiency promotes expansion of proliferative and SHM-active B cells. We next asked what signals might drive the expansion of these DZ-like sub-clusters in the cKO mice. Given our previous work showing αv downregulates the strength of TLR signaling in B cells through action of autophagy proteins [13, 14] we predicted that GC cells that are responsive to TLR signaling could be expanded in cKO mice. Supporting this, clusters 7, 12, 14, and 15 showed elevated expression of *Tlr*s compared to other clusters (**Fig 4F**). Furthermore, lung GC B cells had higher expression of *Tlr*s than equivalent cells in the medLN, providing a potential explanation for the selective effects of αv deletion on lung B cells (**Fig 4G**). Intriguingly, Cluster 13, a DZ-like cluster which also showed higher expression of *Tlr*s, was only found in the medLN of the control mice. One of the highly expressed genes in this cluster was the autophagy gene *Map1lC3b,* and further analysis of GC-related autophagy genes, generated in our previous study [14], showed higher expression of two autophagy-related genes *Map1lC3b* and *Bnip3* in this cluster (**Supp Fig 5F-G**). We have previously shown both *Map1lC3b* and *Bnip3* to be involved in downregulation of TLR signaling[13, 14] therefore, higher expression of these autophagy related genes in Cluster 13 likely suppresses TLR signaling despite elevated receptor expression, explaining why this cluster is not expanded in cKO lungs. These findings suggest that unchecked TLR signaling in αv-deficient B cells likely promotes selective expansion of TLR expressing DZ clusters in cKO mice.

To assess whether αv deficiency altered clonal dynamics, we also analyzed BCR sequences from GC B cells. Analysis of the Inverse Simpson Index, a metric of clonal diversity revealed that lung GC B cells from cKO mice had a lower Inverse Simpson Index than cells from controls, suggesting that cKO lung GC cells undergo greater clonal expansion (**Fig 4H**). To further confirm this effect on clonal expansion, we identified expanded clones (defined as lineages with ≥3 cells sharing a common ancestor) in cKO and control mice. After controlling for differences in the total number of GC B cells in the two genotypes, cKO mice consistently had a higher frequency of expanded clones in both lung and medLN, with the largest differences in lung (**Fig 4I**). Mapping clonal connections to GC subclusters showed that majority of expanded clones in lungs of cKO lungs were related to DZ clusters 7, 12, and 15 (**Fig 4J**), but that expanded clones contributed cells to multiple clusters, linking the DZ subclusters with clonal expansion of GC cells in the lungs of the cKO mice.

Next, we asked whether the expansion of clones in cKO mouse lungs resulted in increased diversification of antigen specificity. PR8-specific GC cells were present in both lung and medLN of all mice (**Fig 4K**). However, cross-reactive GC cells (Cal09-HA^+^ or multiple positive: PR8-HA^+^/Cal09-HA^+^) were present at a much higher frequency in cKO mice than controls. PR8-HA^+^/Cal09-HA^+^ double-positive cells contributed around 5% of HA-specific cells in both lung and medLN of cKO mice but less than 1% of cells in control mice. This difference was even more pronounced for Cal09-HA^+^ single positive cells, which made up almost 25% of GC B cells in cKO mice but were completely absent in lungs of control mice (**Fig 4K-M**). Comparative analysis of SHM in HA+ GC cells showed that both medLN and lung GC cells showed evidence of SHM across genotypes (**Fig 4N**). While the overall mutation burden was similar between genotypes, the cKO lung GC cells exhibited a modest increase in the number of amino acid-replacement mutations per clone (Fig. 4O), suggesting more extensive antigen-driven diversification in the lungs of cKO mice

Collectively, these results indicate that lung GC B cells represent canonical GC B cells and αv integrin restrains the expansion of TLR-responsive DZ-like GC B cells in the lung. In the cKO mice, the loss of this regulation promotes clonal expansion and diversification of antigen specificity, leading to emergence of cross-reactive GC B cells, uniquely within the lungs.

### Loss of B cell αv induces unique memory B cell subsets in the lungs

In contrast to GC B cells, which showed transcriptionally similar populations in lung and medLN, memory B cells showed distinct subclusters in the lung and medLN, based on gene expression (**Fig 5A–B; Supp Fig 5H-I**). In general, memory B cells from medLN displayed higher expression of genes associated with BCR signaling, antigen presentation, and apoptosis, whereas lung memory B cells were enriched for transcripts involved in cell-cycle progression, as well as NF-κB, MAPK, and TLR signaling (**Supp** Fig. 5I).

cKO mice showed striking differences in the relative abundance of memory cell subclusters in the lung compared with controls (**Fig 5A–B**). Cluster 0, the predominant memory cell type in the lungs of control mice, expressed high levels of tissue-resident memory (TRM) B cell markers *Cd69* and *Cd80* (Fig. 5C). In contrast, cluster 1, the predominant memory B cell population in lungs of cKO mice, expressed both canonical memory markers *Fcrl5* and *Cxcr3*, along with high expression of *Cd40* (Fig. 5C**–D**). This cluster also exhibited elevated expression of genes involved in actin cytoskeleton remodeling *Actb* and *Tmsb10* (Fig. 5C), suggesting a distinct migratory or activation state. As with GC cells, lung memory B cells also showed increased expression of TLRs (Fig. 5E), and cluster 1 in particular had elevated expression of *Tlr7* and *Tlr1* (Fig. 5F). These data suggest that αv deficiency promotes expansion of unique memory B cells in the lung.

We next investigated the clonal relationships among HA-specific lung and medLN memory B cells. Lung memory B cells showed substantial clonal expansion in both control and cKO mice, and single clones were represented across different memory subclusters (Fig. 5G). suggesting these clusters represent different activation or differentiation states rather than distinct lineages. Lung memory cells from cKO mice also showed lower sequence diversity than controls, indicating increased clonal expansion (**Fig 5H**), similar to our findings in GC B cells. In contrast, medLN memory B cells showed little to no clonal expansion in either genotype (Fig. 5G), suggesting little local proliferation of these cells. Most HA-specific lung memory B cells were specific for PR8, but a subset of cross-reactive (PR8/Cal09 double-positive or Cal09-specific) memory B cells were detected exclusively in the lungs (Fig. 5I). MedLN-derived memory cells were exclusively PR8-specific and displayed lower levels of somatic hypermutation (SHM) compared to lung-derived counterparts (**Fig 5I–J**). While small memory cell numbers limited statistical comparisons of SHM between genotypes, cKO lung memory cells showed higher total and amino acid replacement mutations (Fig. 5K), consistent with loss of B cell αv integrin facilitating local expansion and diversification of memory B cells in the lung.

Finally, we used BCR sequences to determine the relationship between HA-specific cells in the medLN and lung, and how this was affected by deletion of αv. Majority of lung memory B cells shared sequences with medLN GC cells, consistent with initial activation and seeding from the lymph node (**Supp Fig 5J-K**). A subset of lung GC cells also shared sequences with medLN GCs. However, we also identified GC clones in the lung that did not share sequences with cells in the medLN, raising the possibility that some lung GC cells can develop independently of the medLN (**Supp Fig 5L**). These data, together with the evidence of local clonal expansion (Fig. 4I**, 5G**) support a model in which regardless of the tissue site where the clones originate, a subset of GC and memory B cells expands and evolves within the lung itself. This local evolution is enhanced in the absence of αv integrin, likely through increased responsiveness to innate cues such as TLR signaling, and promotes the emergence of mutated, cross-reactive, GC and memory B cells at mucosal sites.

### Loss of B cell αv promotes sustained increase in lung-resident GC and memory B cells that are cross-reactive and of IgA isotype

The expansion of lung-resident memory B cell subsets in the cKO mice suggested that these mice may exhibit stronger sustained anti-influenza B cell responses than controls. To test this, we measured lung-resident GC and memory B cell responses at day 35 post PR8 IAV infection (**Fig 6A**). By this time, lung-resident GC B cell frequencies and total numbers had returned to baseline levels in control mice, but persisted in cKO mice, comprising up to 4% of IgD^-^ lung-resident B cells (**Fig 6B-C**). Notably, a small subset of the lung-resident GC B cells in cKO mice expressed IgA, which was not seen in controls (**Fig 6C**). Similarly, lung-resident memory B cells persisted at significantly higher levels in cKO lungs compared to controls, with a marked increase in IgA^+^ memory B cells, whereas these had returned to baseline in controls (Fig. 6D). Plasma cell frequencies were low in both genotypes at this time point, although cKO mice showed a modest increase in IgA+ plasma cells (Fig. 6E). Analysis of these subpopulations over time showed a specific increase in lung-resident GC and memory B cells in the cKO mice. Moreover, while IgA^+^ memory B cells comprised a small portion of lung-resident memory B cells, we observed a significant increase in this population over time in the cKO lungs while there was a minimal increase in this population in the control lungs around day 14. In contrast, there was no significant increases in IgM^+^ memory B cells or total plasma cells in the cKO lungs over time indicating specific effects of αv deletion on sustained increase in GC and memory B cells (**Fig 6F**).

We next evaluated the antigen specificity of these lung-resident B cells (**Supp Fig 6A**). PR8-HA-specific and Cal09-HA-reactive GC B cells were still detectable in cKO mice at day 35 but absent in control mice (Fig. 6G). Antigen-specific memory B cells were rare overall, with PR8-specific memory cells found only in cKO lungs (**Supp Fig 6B**) while frequencies of Cal09-reactive memory were very low in both genotypes (**Supp Fig 6C**).

To further characterize these small numbers of antigen-specific cells that were expanded in the cKO mice, we performed high-dimensional clustering analysis of lung-resident IgD⁻ HA-specific B cells using PhenoGraph (Cytofkit, Bioconductor). This analysis identified 11 statistically distinct clusters (Fig. 6H). We found 4 of these clusters (11, 4, 2 and 7) had a higher frequency of cells in the cKO mice, compared to controls and we assessed the characteristics of these expanded cells (**Fig 6I**). All of these 4 clusters showed higher IgA expression and increased staining for Cal09-HA, compared to other clusters (**Fig 6J**). Moreover, these 4 clusters also showed increased expression of GC-associated markers (Fas or PNA), compared to other clusters (**Fig 6J**). In contrast, the clusters that had a higher frequency in the control mice compared to the cKO (cluster 6, 9, and 8) showed increased PR8-HA staining and IRF4 expression (**Fig 6J**) indicating increase in plasma cells recognizing infecting virus. This analysis further confirmed that the loss of αv promotes the persistence of IgA-expressing, cross-reactive GC B cells in the cKO lung.

To confirm the antigen reactivity of the memory B cells that were expanded in the cKO lungs, we performed a similar cluster analysis on memory B cells generated at day 4 post re-challenge, when the memory response would be at peak and provide more cells for analysis of this compartment. We also used CD73 as a positive marker for memory B cells, based on our scRNAseq analysis, where we found expression of CD73 (Nt5e) on memory subclusters in the lung. We performed PhenoGraph cluster analysis within the lung-resident IgD^-^ memory B cells (**Fig 6K**) and identified four clusters (8, 1, 6, and 7) that were more frequent in cKO mice (**Fig 6L**). These 4 clusters were highly positive for Cal09-HA and the marker CD73 (**Fig 6M**). There was only one cluster showing high IgA expression, cluster 8 (**Fig 6M**). This was one of the clusters with increased frequency of cells in the cKO mice and showed high CD73 expression as well as increased staining for Cal09-HA (**Fig 6M**). CD73 is a marker associated with GC derived memory B cells[25], therefore these findings indicate that the loss of B cell αv leads to sustained increase in GC B cells in the lungs after infection, which leads to increases in cross-reactive IgA^+^ memory B cells.

### Persistence of GC reactions due to loss of B cell αv leads to the sustained increase in cross-reactive GC, memory and IgA+ B cells

To confirm that the observed increases in lung-resident IgA^+^ GC and memory B cells in cKO mice are due to ongoing germinal center (GC) activity in iBALTs we analyzed lungs of cKO and control mice at day 36 post infection by confocal microscopy.

At day 36 post infection, cKO mice had clearly identifiable GC clusters in iBALT areas, consistent with the detection of GC B cells by FACs. Curiously, we also detected areas of GL7-staining cells in control mice, and these were of a similar size to those in cKO mice (**Fig 7A**). This was at odds with our FACs analysis in which we detected fewer GC B cells in control mice. However, this apparent discrepancy may be due to the much lower staining for GL7 in the control mice (**Fig 7A-C**). Notably, cKO mice showed a large increase in IgA^+^ cells in the lung compared with controls (**Fig 7A**). Quantification confirmed that both the total area of IgA^+^ cells (**Fig 7D**) and the percentage of IgA^+^ GL7^+^ (**Fig 7E**), within iBALT-GC structures remained significantly elevated in cKO mice. Moreover, in the cKO lungs we saw significant amount of IgA^+^ cells also outside the GC areas which we interpret as the lung-resident IgA^+^ memory B cells (**Fig 7F**) observed in the flow cytometry analysis. We also observed Cal09-HA+ cells within GC regions of cKO lungs, but these were not detected in controls (**Fig 7G-H**), further supporting the presence of cross-reactive GC B cells locally in the lungs of the cKO mice

**Figure 7.**
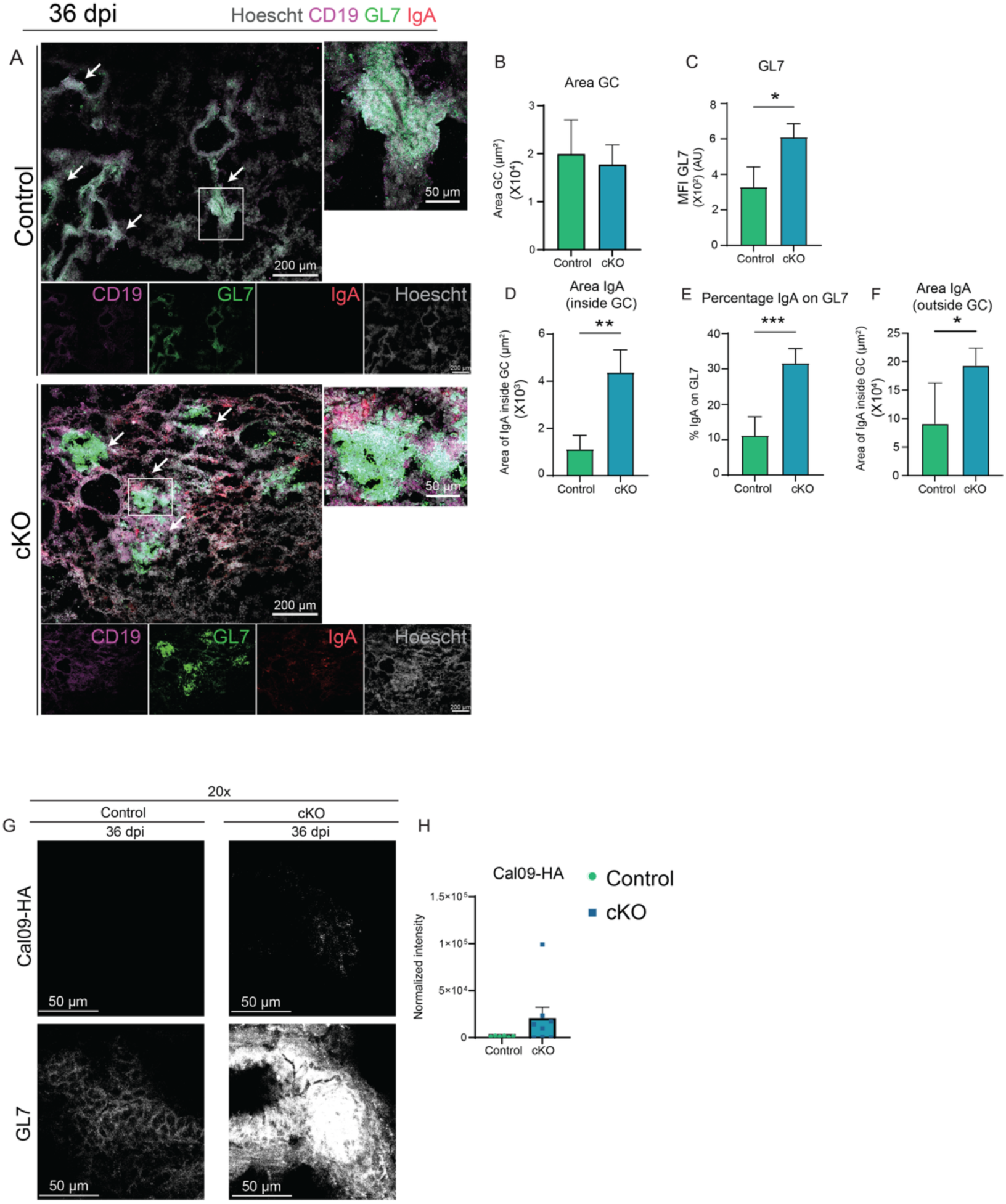
Absence of αv B cell integrin leads to persistence of GC reactions in the lungs after influenza infection. αv^+/+^ CD19^Cre+^ (control) and αv^fl/fl^ CD19^Cre+^ (cKO) mice were infected i.n. with 100 plaque-forming units (PFU) of live PR8 IAV. Lungs were harvested after 36 days post infection (dpi) (**A**) Representative confocal images of lung sections of one single plane from infected control (top) and cKO mice (bottom) at 20x magnification, scale bar 200 µm. Lung staining for Hoescht (grey), CD19 (magenta), GL7 (green) and IgA (red). Arrows represent GL7 positive structures corresponding to iBALTs. The insets highlight iBALTs, scale bar 50 µm. (**B – F**) Bar graph showing mean and SEM of the GC area in µm^2^ (**B**), GL7 mean fluorescence intensity (MFI) within the iBALT structures (**C**), IgA area inside the GC in µm^2^ (**D**), percentage of IgA staining overlapping with GL7 within the iBALT (**E**), and IgA area outside the GC structures (**F**). Quantifications made from 9-12 different fields of sections viewed at 10x from 3 different mice per genotype (N= 12-27). (**G**) Representative confocal images of iBALT regions, stained for Cal09-HA (top) and GL7 (bottom) of lungs from infected control (left) and cKO (right) mice at 20x magnification. (**H**) Bar graph of the normalized intensity of Cal09-HA staining from 6 – 8 different fields of sections viewed at 20x from 3 different mice per genotype. **p<0.01, ***p<0.001 by Mann-Whitney U-test.

Together, these findings demonstrate that B cell-specific deletion of αv integrin results in prolonged ectopic GC activity in the lungs which contributes to the local generation and maintenance of GC and memory B cells cross-reactive to antigenic variants of the infecting virus and of the IgA isotype.

### B cell αv integrin regulates IgA differentiation in a B cell intrinsic manner

Our scRNA-seq analysis revealed that the expansion of germinal center (GC) B cells in the lungs of αv-deficient (cKO) mice was associated with elevated expression of Toll-like receptors (TLRs), particularly within DZ-like subclusters. Consistent with this, we previously showed that αv integrin restrains TLR signaling in systemic immune responses, limiting TLR-induced class switching to IgG2c. However, it is unclear whether increased TLR signaling could be responsible for the expansion of IgA class-switched cells that we see in lungs of cKO mice. To test this, we used *in vitro* models of B cell differentiation and class switching.

We first tested whether TLR stimulation could promote greater amounts of IgA class switching in the absence of αv. B cells isolated from either lung or spleen were stimulated with the R848, a synthetic ligand for TLR7, and IgA class switching was measured by flow cytometry. R848 was sufficient to promote generation of IgA^+^ B cells in the lung B cell culture, and approximately twice as many IgA^+^ B cells were generated from lungs of cKO mice than controls (**Fig 8A**), in agreement with our model. In contrast B cells from the spleen, regardless of the genotype, did not differentiate into IgA cells under these conditions (**Fig 8B**). We speculated that this may reflect the lack of exposure of spleen B cells to the mucosal conditioning signals such as retinoic acid and TGF-β, that promote IgA class switching. To more comprehensively model the full process of IgA class-switching and assess the contribution of TLR signaling, we turned to a multi-step culture protocol that better models all aspects of B cell activation and IgA differentiation[26, 27]. Total IgA⁻ lung-resident B cells from lung were first activated via TLRs using CpG (a TLR9 ligand) to engage both naïve and activated B cells, followed by co-stimulation with BAFF and CD40L to mimic germinal center-like conditions. IL-21, retinoic acid, and TGF-β were also included to promote IgA class switching (**Fig 8C**). Under these conditions, IgA^+^ B cells were generated in all cultures, at much higher frequency than in cells treated with R848 alone. However, cKO B cells consistently generated higher proportions of IgA^+^ B cells compared with control B cells (**Fig 8D**). In parallel, significantly more IgA was detected in the supernatants of cell cultures from cKO mice (**Fig 8E**). Cultures of B cells isolated from spleen also led to class switch to IgA isotype (**Fig 8F**) and the spleen B cell culture from cKO mice showed increase in IgA^+^ cells. However, the fold-increase in IgA^+^ cells was significantly greater in lung-derived cKO B cells than in spleen-derived cells (Fig. 8G), suggesting mucosal imprinting enhances TLR-driven IgA differentiation.

**Figure 8.**
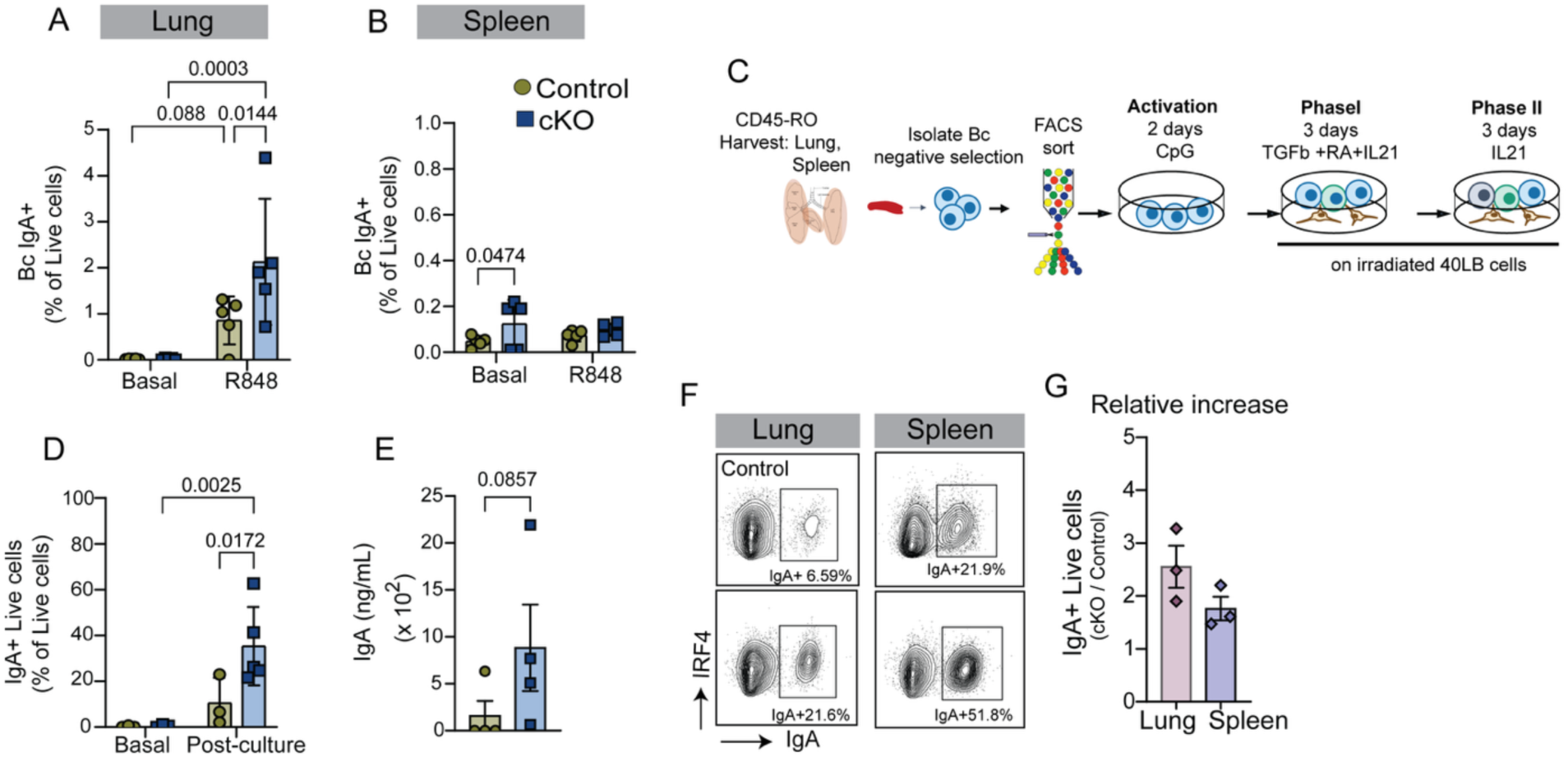
TLR-ligand stimulation leads to the differentiation and proliferation of tissue resident IgA-class switched B cells. **(A-B)** Lung and spleen B cells were isolated from uninfected control and cKO mice and cultivated in a 96 well plate with R848 (1 μg/mL) for 4 days. IgA expression was assessed by flow cytometry. Quantification of the frequency of IgA-expressing B cells in the lungs **(A)** and spleen **(B)** of control and cKO mice. (**C**) Schematic of the experimental procedure. C57BL/6 (control) and αv^fl/fl^ CD19^Cre+^ (cKO) mice were injected r.o. with 1µg α-CD45-APC five minutes prior to euthanasia. Lung and spleen were then collected and sorted for i.v.CD45-unlabeled resident cells and plated for a total of 8 days. Cells were activated with CpG (1 μg/mL) for 2 days and then added TGFβ (2ng/mL), Retinoic acid (100nM) and IL-21 (40 ng/mL). (**D**) Frequency of IgA-expressing live cells within lung-resident B cells after culture as determined by flow cytometry. (**E**) Secreted IgA as measured by ELISA of the supernatant of lung-resident B cells after TGFβ, RA and IL-21 culture. (**F**) Representative flow plots of lung and spleen resident B cells after the entire 8-day culture protocol. Cells were gated from live lymphocytes. (**G**) Relative increase in the frequency of IgA+ live cells, in the cKO vs the control cultures comparing cultures from lung and spleen B cells after the 8-day culture protocol. Each dot represents culture with cells from individual mice from 3-5 independent experiments. Bar graphs show mean and SEM±, statistical significance determined by 2-way ANOVA (**A**, **B** and **D**) or Mann-Whitney U-test (**E**).

Together, these findings demonstrate that B cell–intrinsic αv integrin limits TLR-mediated class switching to IgA, particularly in mucosally conditioned lung B cells. Loss of αv enhances the responsiveness of these cells to TLR signals, resulting in increased IgA class switching *in vitro.* This is consistent with our *in vivo* findings of sustained increase in IgA^+^ GC and memory B cell responses in the lungs of cKO mice. These results support a model in which αv integrin functions as a key checkpoint that restricts mucosal B cell activation and class-switching by restraining GC activity of TLR-responsive B cells following respiratory viral infection.

## Discussion

Recent studies have shown that natural infection of the respiratory tract by influenza virus induces memory B cells that are lung-resident[2], major producers of broadly-reactive antibodies[28] and IgA isotype antibodies[2, 4]. These tissue-resident B cells contribute substantially to secondary protection but are not efficiently generated by systemic vaccines, highlighting a critical need to understand the local cues that shape mucosal B cell immunity. Here, we identify B cell αv integrin as a key negative regulator of GC persistence, clonal evolution, and IgA+ memory B cell development in the lung following influenza virus infection. Loss of B cell αv promoted expansion of GC and memory B cells in the lung and generation of cross-reactive B cells in the lungs compared to the medLN, revealing a spatially restricted mechanism of mucosal B cell regulation.

GCs are sites of affinity maturation of B cells, where B cells undergo successive rounds of proliferation, SHM and selection, through antigen presentation to T follicular helper (Tfh) cells[29]. Studies on influenza and other viral infections have revealed that increasing SHM in the GCs not only increases the affinity of antibodies for immunizing antigens but also their breadth, enabling recognition of diverse viral strains[30, 31]. While we previously showed that αv loss enhances GC responses and SHM in secondary lymphoid organs, its role in mucosal B cell responses was unknown. Here, we show that αv-deficient B cells form persistent, ectopic GCs within inducible bronchus-associated lymphoid tissue (iBALT) in the lungs (Fig 7), leading to expanded clones with elevated SHM and increased cross-reactivity to heterologous HA variants (Fig 4 **and 5**). These structures persist for at least 35 days post-infection and support the formation of IgA^+^ memory B cells in the respiratory mucosa.

Using single-cell transcriptomics of antigen-specific B cells, we observed that influenza infection induces GC and memory B cells with shared clonal origins in the lung and medLN. However, in αv-deficient mice, we observed preferential expansion of DZ-like GC subclusters in the lung enriched for TLR expression (Fig 4), which correlated with increased SHM and cross-reactivity (Fig 4). Lung-resident memory B cells in these mice also exhibited greater clonal expansion and broader HA specificity than their medLN counterparts (Fig 5), indicating that prolonged GC activity at mucosal sites shapes the local memory B cell pool. These findings align with previous studies showing that lung-resident memory B cells are key contributors to cross-reactivity[28] and underscore that local B cell evolution driven by tissue-intrinsic cues, likely involving enhanced innate immune signaling are key to promoting expansion of cross-reactive B cells in the lungs

We have previously shown that the loss of B cell αv enhances responses from innate-like B cell populations such as B1 B cells, Marginal zone B cells, as well as extrafollicular B cells[6, 15]. These B cell populations can all be involved in the immune response against respiratory viruses[1, 32] and likely lead to IgA production at the respiratory tract associated with early clearance of the virus. But here we focused on lung-resident GC-derived B cells that emerge later and are more relevant for secondary protection. In αv cKO mice, persistent iBALT structures supported accumulation of IgA^+^ GC and memory B cells from day 16 through day 36 post-infection (Fig 3 **and 7**). Upon secondary challenge, cKO lungs harbored increased numbers of CD73+ IgA+ and Cal09-binding memory B cells (Fig 6). While we cannot rule out the possibility that these memory B cells could be generated from GC reactions in the medLN, based on our RNAseq data we predict that the increases in cross-reactive IgA memory B cells in the lungs of cKO mice are at least in part due to increased expansion of GC and memory B cells in the lungs.

Although IgA^+^ cells were a minor fraction of the lung-resident memory B cell pool, their selective enrichment in the cKO mice underscores a previously unrecognized opportunity to drive GC-derived long-lived mucosal IgA immunity. While IgA^+^ cells were underrepresented in our scRNA-seq dataset, likely due to low abundance, transcriptomic analysis revealed distinct lung memory B cell subsets enriched for TLR, NF-κB, and cytoskeletal remodeling pathways-signatures associated with heightened activation and tissue residency. We predict that these transcriptional programs are linked to the expanded IgA^+^ memory population in cKO lungs and arise via TLR-driven signals. Further studies at later time points will be necessary to confirm this relationship. Importantly, a major barrier to studying lung-resident memory B cells is the lack of specific surface markers for their isolation. Our scRNA-seq data provide transcriptional signatures that distinguish lung and medLN memory subsets, which will enable functional dissection of the protective capacity of lung-resident memory cells in future. Moreover, while further studies are still necessary to establish the full heterosubtypic protective capacity of lung-resident GC and memory responses against multiple influenza strains our studies provide a basis to design these studies.

Mechanistically, our *in vitro* experiments demonstrated that B cell-intrinsic loss of αv enhanced IgA class switching in response to TLR stimulation, especially under conditions mimicking mucosal cytokine environments (Fig 8). This aligns with prior findings that αv deficiency leads to enhanced TLR-induced expression of *Aicda*[14, 16], a critical gene for class-switch recombination and SHM. Notably, the increase in IgA class switching was more pronounced in B cells derived from the lung than from the spleen, further supporting a tissue-specific effect. While αv loss also increases IgG responses, we focused here on IgA^+^ B cells, as their developmental regulation in respiratory tissues is less well understood. TGFβ-driven IgA class switching has been described in the gut and IgA B cells are known to receive stronger positive selection cues in Peyer’s Patch GCs[33], but the mechanisms governing lung IgA B cells remain poorly defined. Our findings suggest that TLR signals-amplified in the absence of αv-support IgA class switching in lung B cells, uncovering a novel mucosal regulatory axis.

Our findings highlight B cell αv integrin as a novel checkpoint that constrains mucosal GC activity and IgA^+^ memory B cell diversification in the respiratory tract. These results have important implications for vaccine development. Given growing evidence that mucosal IgA responses limit infection and transmission of respiratory viruses more effectively than systemic IgG responses[4, 34] [35, 36], these findings offer a promising strategy for improving respiratory virus vaccines. A key challenge with current respiratory viral vaccines is the lack of protection from infection at respiratory sites as well as the lack of durability of protection induced by the vaccines. Strategies that prolong GC activity in the lung through intranasal vaccines could drive both protection at respiratory sites and promote long-lived GC driven IgA^+^ memory B cell responses. Intranasal vaccines incorporating αv blockade or TLR ligand adjuvants may be particularly effective in this regard. Future studies on pharmacological αv integrin blockade and its overall consequences would be valuable to assess the translation capacity of these findings. Moreover, studies on the effects of TLR ligand adjuvants in mucosal vaccines could similarly be used to specifically promote lung-resident long-lived B cell responses for vaccine efficacy.

## Materials and Methods

### Mice

αv^fl/fl^ mice were backcrossed to C57BL/6J mice as previously described[13, 16]. The animals were bred and maintained on the C57BL/6 background. αv^fl/fl^ mice were crossed with CD19 cre transgenic mice to generate B cell specific αv knockout mice, referred to as conditional KO (cKO) mice (CD19^cre/+^ αv^fl/fl^). Mice with a single CD19^Cre^ allele, CD19^cre/+^αv ^+/+^ were used as controls. For *in vitro* experiments C57BL/6 mice were used as controls. All mice for this study were age and sex-matched and were 9-14 weeks old at the time of infectio. Mice were housed in a specific pathogen-free facility. All procedures were approved by the Institutional Animal Care and Use Committee at Seattle Children’s Research Institute.

### Influenza infection

Live PR8 IAV was purchased from Charles River Laboratories and stored in single-use aliquots at −80°C. Just prior to infection, PR8 was thawed and diluted in PBS. Mice were anesthetized with Isoflurane (Patterson Veterinary) and infected with PR8 IAV in 25µL in the nostrils. Mice were weighed and monitored for signs and symptoms of disease for the duration of the study.

### HA tetramer preparation

The B cell HA tetramers were either a gift from Dr. Emily Gage[17] or from Dr. Kanekiyo Masaru’s lab at NIH. Tetramers with an AviTag were biotinylated with the BirA biotin-ligase kit (Avidity) according to manufacturer’s instructions and stored in single-use aliquots at −80°C. Following biotinylation, BV711-(BioLegend) or PerCpCy5.5-(BioLegend) conjugated streptavidin or TotalSeq-SA-C0951-PE (BioLegend) was added in 5 serial steps with a 10-minute incubation period each to the unlabeled tetramers. Labeled tetramers were prepared the day before the experiment.

### Antibodies

α-IgD-BV786, α-B220-BUV395, α-CD45-BUV395, α-CD19-BUV737 were purchased from BD Horizon. Α-CD3-BV510, α-CD11c-BV510, α-F4/80-BV510, α-CD138-BV421, α-Fas-BV605, Streptavidin (SA)-PerCPCy5.5, SA-BV711, α-IgM-PerCPCy5.5 and TotalSeq-C0301 – C0315 and TotalSeq-SA-C0951 (for scRNAseq) were from BioLegend. α-CD45-APC, α-IRF4-PeCy7 and α-IRF4-AF700 were purchased from Invitrogen. α-IgA-Biotin, α-IgA-PE and α-IgM-PeCy7 were from Southern Biotech. α-Gr1-BV510 and PNA-FITC were from Miltenyi Biotechnologies and Vector Labs, respectively.

### Flow cytometry

5 minutes prior to euthanasia, mice were injected with 1µg α-CD45-BUV395 or α-CD45-APC in the retro-orbital cavity. Lungs and mediastinal lymph nodes (medLN) were collected from euthanized mice. Lung tissue was digested with 1.5mg/mL collagenase (Milipore Sigma) and 10µg/mL DNAse I (Milipore Sigma) in Mg^2+^, Ca^2+^ free Hank’s balanced salt solution (HBSS, Gibco) with 10% fetal bovine serum (FBS, Sigma) for 1 hr at 37°C or 30 min in a shaking incubator at 1000 rpm. Lungs and medLN were then ground into a 40µm filter to generate a single cell suspension. For some experiments lung B cells were isolated using CD19 positive magnetic bead separation (STEMCell). Cells were counted using AccuCheck count beads (ThermoFisher) or Muse Cell Analyzer (Luminex) according to the manufacturer’s instructions. Cells were then stained with fixable LiveDead Aqua (ThermoFisher) to identify dead cells, after which cells were stained with PR8-HA and Cal09-HA B cell tetramers and surface antibodies for 20 minutes at room temperature. To identify intracellular antigens, cells were then fixed and permeabilized with the Transcription Factor kit (BioLegend) according to manufacturer’s instructions. Events were analyzed on an LSRII flow cytometer (BD Biosciences) or an LSR Fortessa (BD Biosciences) and analyzed using FlowJo (v.10, TreeStar). See Supplemental Figure 1 for gating strategies.

Cells for sorting were prepared as a single cell suspension (as described above) from lungs and medLN of PR8 IAV infected mice. 5 minutes prior to euthanasia, mice were injected with 1µg α-CD45-APC in the retro-orbital cavity. Prior to staining, cells were blocked with Fc block for 10 minutes and then stained with fixable LiveDead Aqua (ThermoFisher) to identify dead cells, surface antibodies, hashtag oligos for each mouse and PR8-HA and Cal09-HA B cell tetramers for 20 minutes at 4C. Cal09-HA and PR8-HA tetramers were bound to SA-Oligo tagged-PE and SA-BV711 to gate out SA-PE specific B cells from the lungs and medLN. All cells from each mouse were combined into a single pool per tissue and sorted to enrich for tetramer-specific cells gated as live *CD45-unlabeled B220^+^ HA-Tetramer++*.

### ELISpot

ELISpots were performed on single cell suspensions of lung or medLN cells, prepared as described above. MAIPS ELISpot plates (Millipore) were pre-coated with 3µg/mL PR8-HA or Cal09-HA (SinoBiological) diluted in PBS. Serial dilutions of cells were plated over wells and incubated over night at 37°C. Spots were developed using α-mouse IgG or α-mouse IgA alkaline phosphatase (AP) Abs (Southern Biotech) followed by BCIP/NBT AP substrate (Vectors Labs). Spots were imaged and counted using an ImmunoSpot analyzer (Cellular Technology Limited).

### Histology and Immunofluorescence

Lungs were perfused with PBS and then inflated and embedded in OCT, and frozen at −80°C (as described in[37]). Frozen sections (8μm) were fixed in acetone at −20°C, air dried and blocked with 1% normal goat serum (Jacskon ImmunoResearch) and 1% BSA (Sigma) in PBS with 0.1% Tween-20 (Sigma). Slides were stained with Hoescht, α-CD19-AF647, α-GL7-FITC and α-IgA-PE. Images were acquired with a Zeiss LSM 900 confocal microscope. Image processing and analysis were performed with FIJI (ImageJ) software[38], with a personalized macro where the iBALT structure was manually delimited by the polygon tool, and then the image was converted to a mask by default threshold to measure the area of IgA and GL7 positive staining by the “analyze particle” function. IgA+GL7+ positive area percentage was calculated by applying the image Calculator “AND” function over IgA and GL7 masks, then measuring the resulting area by analyze particle function and calculating the percentage of the resulting area over the whole GL7 positive area.

### Single cell RNA sequencing

B Cells were isolated from lung and medLN from mice at 20 days post infection with PR8 influenza (as described above). Enriched HA-tetramer specific B cells were run on 10x Chromium with 5’ GEM-X kit to isolate single cells, and libraries generated for gene expression, BCR sequencing, and hashtag oligo quantification. Libraries were sequenced on an Illumina NextSeq 2000 following 10x guidelines for read formats. Libraries were processed to gene and hashtag UMI counts, and assembled BCR sequences with isotype, using cellranger v. 8.0.0. Downstream analysis was performed using Seurat v. 5.1 [39]. Cells were filtered for quality using standard metrics, expression of target and non-target cell type genes, then demultiplexed using hashtag oligo counts. Cells were further filtered based on detection of at most one heavy and two light BCR chains, yielding 9660 high-quality single B cells. Normalization, UMAP projection, clustering, and marker gene identification were performed with Seurat, and clusters were assigned to major B cell subsets using marker genes *Ighd, Aicda, Cd80* and *Prdm1*. Cells from mice lacking substantial GC B cell populations in the medLN were excluded from downstream analysis. Cells were called positive or negative for each HA tetramer using a threshold on log-transformed barcode UMI counts. We used SCENIC to identify modules of genes putatively regulated by transcription factors[40].

Analysis of BCR sequences was conducted using the Immcantation toolkit. We used Change-O to assign sequences to V and J genes, define clones by matched using heavy and light chains genes and an empirical threshold on CDR3 Hamming distances, infer germline sequences [41]. Trees were inferred using the dowser implementation of IgPhyML and the HLP19 model on paired heavy and light chains [42–44]. Somatic hypermutation was quantified using SHazaM [41]. Clonal diversity metrics were calculated by repeatedly down sampling to a common number of cells per mouse in the focal population, then calculating the metric on each down sampled replicate.

### In vitro plasma cell culture

All mice were injected retro orbitally with 1µg αCD45-BUV395 5 minutes prior to euthanasia. B cells from spleen, medLN or lung were isolated using negative magnetic bead separation (STEMCell). Cultures with R848 only were performed on lympholyte-M (Cedarlane) separated lung B cells and B cells from spleen isolated using negative magnetic bead separation. B220^+^ B cells were then sorted for i.v. CD45^-^ populations using an Aria FACS sorter (BD Biosciences). Plasma cells were differentiated from naive B cells. Briefly, cells were suspended in Iscove’s modified Dulbecco’s medium (IMDM, cytiva) supplemented with 10% FBS (Sigma), 2mM Glutamax (Gibco), 55μM β-mercaptoethanol (Gibco), 100U/mL penicillin (Gibco) and 100μg/mL streptomycin (Gibco) and activated with 1µg/mL CpG (Invivogen) for two days. Cells were then seeded over irradiated 40LB feeder cells, a generous gift from Dr. Daisuke Kitamura[27], expressing mouse CD40L and BAFF. To induce class switch to IgA, cells were incubated with 2ng/mL TGF-β (BioLegend), 100nM retinoic acid (RA, Sigma) and 40ng/mL IL-21 (Peprotech) for three days, followed by three days in IL-21 alone.

#### ELISA

Enzyme linked immunosorbent assays (ELISAs) were performed on the supernatant of cultured cells (see *in vitro* plasma cell culture for culture details). Immulon 2HB plates (ThermoFisher) were pre-coated with 2μg/mL of goat-α-mouse Ig(H+L) (Southern Biotech) diluted in PBS. Serial dilutions of supernatant were plated and left overnight at 4°C. The plates were developed using α-mouse IgA-AP Abs (Southern Biotech) followed by 4-Nitropenyl phosphate disodium salt hexahydrate substrate tablets (Sigma). Absorbance was measured using a Spectramax 190 luminometer (Molecular Devices) at 405 nm.

### Statistical analysis

Raw data from control and cKO PR8-infected groups were analyzed by Mann-Whiney U test or 2-way ANOVA using GraphPad Prism (v.9.3.1). *p*<0.05 were considered statistically significant.

## Abbreviations

GC: germinal center
IAV: Influenza A Virus
TLR: Toll-like receptor
SHM: somatic hyper mutation
medLN: Mediastinal Lymph Node
cKO: av conditional knockout mice
HA: Hemagglutinin
iBALT: inducible bronchus-associated lymphoid tissues.

## Acknowledgements

We thank Sara Sagadiev, Aaron Liu and Ursula Holder for laboratory assistance and useful insights and discussion. This research was supported by the Office of Animal Care, the Microscopy and Histopathology CoLab, the Flow Cytometry Core at Seattle Children’s Research Institute and the Genomics Core at the Benaroya Research Institute. We are grateful to Dr. K. Masaru and Dr. R. Gillespie at NIH and Dr. Emily Gage for the influenza tetramers and guidance on tetramer staining.

## References

1. Lam, J.H. and N. Baumgarth, The Multifaceted B Cell Response to Influenza Virus. J Immunol, 2019. 202(2): p. 351–359.

2. Allie, S.R., et al., The establishment of resident memory B cells in the lung requires local antigen encounter. Nat Immunol, 2019. 20(1): p. 97–108.

3. Onodera, T., et al., Memory B cells in the lung participate in protective humoral immune responses to pulmonary influenza virus reinfection. Proc Natl Acad Sci U S A, 2012. 109(7): p. 2485–90.

4. Oh, J.E., et al., Intranasal priming induces local lung-resident B cell populations that secrete protective mucosal antiviral IgA. Sci Immunol, 2021. 6(66): p. eabj5129.

5. Acharya, M., et al., SDF-1 and PDGF enhance alphavbeta5-mediated ERK activation and adhesion-independent growth of human pre-B cell lines. Leukemia, 2009. 23(10): p. 1807–17.

6. Acharya, M., et al., *alphav Integrin expression by DCs is required for Th17 cell differentiation and development of experimental autoimmune encephalomyelitis in mice*. J Clin Invest, 2010. 120(12): p. 4445–52.

7. Gianni, T., et al., alphavbeta3-integrin is a major sensor and activator of innate immunity to herpes simplex virus-1. Proc Natl Acad Sci U S A, 2012. 109(48): p. 19792–7.

8. Lacy-Hulbert, A., et al., Ulcerative colitis and autoimmunity induced by loss of myeloid alphav integrins. Proc Natl Acad Sci U S A, 2007. 104(40): p. 15823–8.

9. Lucas, M., et al., Apoptotic cells and innate immune stimuli combine to regulate macrophage cytokine secretion. J Immunol, 2003. 171(5): p. 2610–5.

10. Overstreet, M.G., et al., Inflammation-induced interstitial migration of effector CD4(+) T cells is dependent on integrin alphaV. Nat Immunol, 2013. 14(9): p. 949–58.

11. Savill, J., et al., Vitronectin receptor-mediated phagocytosis of cells undergoing apoptosis. Nature, 1990. 343(6254): p. 170–3.

12. Schrock, D.C., et al., Pivotal role for alphaV integrins in sustained Tfh support of the germinal center response for long-lived plasma cell generation. Proc Natl Acad Sci U S A, 2019.

13. Acharya, M., et al., *alphav Integrins combine with LC3 and atg5 to regulate Toll-like receptor signalling in B cells*. Nat Commun, 2016. 7: p. 10917.

14. Muir, V., et al., Transcriptomic analysis of pathways associated with ITGAV/alpha(v) integrin-dependent autophagy in human B cells. Autophagy, 2023. 19(3): p. 926–942.

15. Acharya, M., et al., B Cell alphav Integrins Regulate TLR-Driven Autoimmunity. J Immunol, 2020. 205(7): p. 1810–1818.

16. Raso, F., et al., *alphav Integrins regulate germinal center B cell responses through noncanonical autophagy*. J Clin Invest, 2018. 128(9): p. 4163–4178.

17. Gage, E., et al., Memory CD4(+) T cells enhance B-cell responses to drifting influenza immunization. Eur J Immunol, 2019. 49(2): p. 266–276.

18. Matsumoto, R., et al., Induction of bronchus-associated lymphoid tissue is an early life adaptation for promoting human B cell immunity. Nat Immunol, 2023. 24(8): p. 1370–1381.

19. Moyron-Quiroz, J.E., et al., Role of inducible bronchus associated lymphoid tissue (iBALT) in respiratory immunity. Nat Med, 2004. 10(9): p. 927–34.

20. Denton, A.E., et al., Type I interferon induces CXCL13 to support ectopic germinal center formation. J Exp Med, 2019. 216(3): p. 621–637.

21. de Carvalho, R.V.H., et al., Clonal replacement sustains long-lived germinal centers primed by respiratory viruses. Cell, 2023. 186(1): p. 131–146 e13.

22. Kanekiyo, M., et al., Mosaic nanoparticle display of diverse influenza virus hemagglutinins elicits broad B cell responses. Nat Immunol, 2019. 20(3): p. 362–372.

23. Kennedy, D.E., et al., Novel specialized cell state and spatial compartments within the germinal center. Nat Immunol, 2020. 21(6): p. 660–670.

24. Pae, J., et al., Transient silencing of hypermutation preserves B cell affinity during clonal bursting. Nature, 2025. 641(8062): p. 486–494.

25. Conter, L.J., et al., CD73 expression is dynamically regulated in the germinal center and bone marrow plasma cells are diminished in its absence. PLoS One, 2014. 9(3): p. e92009.

26. Hung, K.L., et al., Engineering Protein-Secreting Plasma Cells by Homology-Directed Repair in Primary Human B Cells. Mol Ther, 2018. 26(2): p. 456–467.

27. Nojima, T., et al., In-vitro derived germinal centre B cells differentially generate memory B or plasma cells in vivo. Nat Commun, 2011. 2: p. 465.

28. Adachi, Y., et al., Distinct germinal center selection at local sites shapes memory B cell response to viral escape. J Exp Med, 2015. 212(10): p. 1709–23.

29. Victora, G.D. and M.C. Nussenzweig, Germinal centers. Annu Rev Immunol, 2012. 30: p. 429–57.

30. Corti, D., et al., Cross-neutralization of four paramyxoviruses by a human monoclonal antibody. Nature, 2013. 501(7467): p. 439–43.

31. Pappas, L., et al., Rapid development of broadly influenza neutralizing antibodies through redundant mutations. Nature, 2014. 516(7531): p. 418–22.

32. Baumgarth, N., How specific is too specific? B-cell responses to viral infections reveal the importance of breadth over depth. Immunol Rev, 2013. 255(1): p. 82–94.

33. Raso, F., et al., Antigen receptor signaling and cell death resistance controls intestinal humoral response zonation. Immunity, 2023. 56(10): p. 2373–2387 e8.

34. Kwon, D.I., et al., Mucosal unadjuvanted booster vaccines elicit local IgA responses by conversion of pre-existing immunity in mice. Nat Immunol, 2025. 26(6): p. 908–919.

35. Hassan, A.O., et al., A Single-Dose Intranasal ChAd Vaccine Protects Upper and Lower Respiratory Tracts against SARS-CoV-2. Cell, 2020. 183(1): p. 169–184 e13.

36. Hassan, A.O., et al., An intranasal vaccine durably protects against SARS-CoV-2 variants in mice. Cell Rep, 2021. 36(4): p. 109452.

37. Davenport, M.L., et al., Perfusion and Inflation of the Mouse Lung for Tumor Histology. J Vis Exp, 2020(162).

38. Schindelin, J., et al., Fiji: an open-source platform for biological-image analysis. Nat Methods, 2012. 9(7): p. 676–82.

39. Hao, Y., et al., Dictionary learning for integrative, multimodal and scalable single-cell analysis. Nat Biotechnol, 2024. 42(2): p. 293–304.

40. Aibar, S., et al., SCENIC: single-cell regulatory network inference and clustering. Nat Methods, 2017. 14(11): p. 1083–1086.

41. Gupta, N.T., et al., Change-O: a toolkit for analyzing large-scale B cell immunoglobulin repertoire sequencing data. Bioinformatics, 2015. 31(20): p. 3356–8.

42. Hoehn, K.B., G. Lunter, and O.G. Pybus, A Phylogenetic Codon Substitution Model for Antibody Lineages. Genetics, 2017. 206(1): p. 417–427.

43. Hoehn, K.B., O.G. Pybus, and S.H. Kleinstein, Phylogenetic analysis of migration, differentiation, and class switching in B cells. PLoS Comput Biol, 2022. 18(4): p. e1009885.

44. Hoehn, K.B., et al., Repertoire-wide phylogenetic models of B cell molecular evolution reveal evolutionary signatures of aging and vaccination. Proc Natl Acad Sci U S A, 2019. 116(45): p. 22664–22672.

