## Supplemental file for "B cell αv integrin regulates tissue specialization and clonal expansion of lung germinal center and memory B cells after viral infection"

[*]Corresponding author:

**This pdf file includes:**

Supplementary text

Figures 1-6

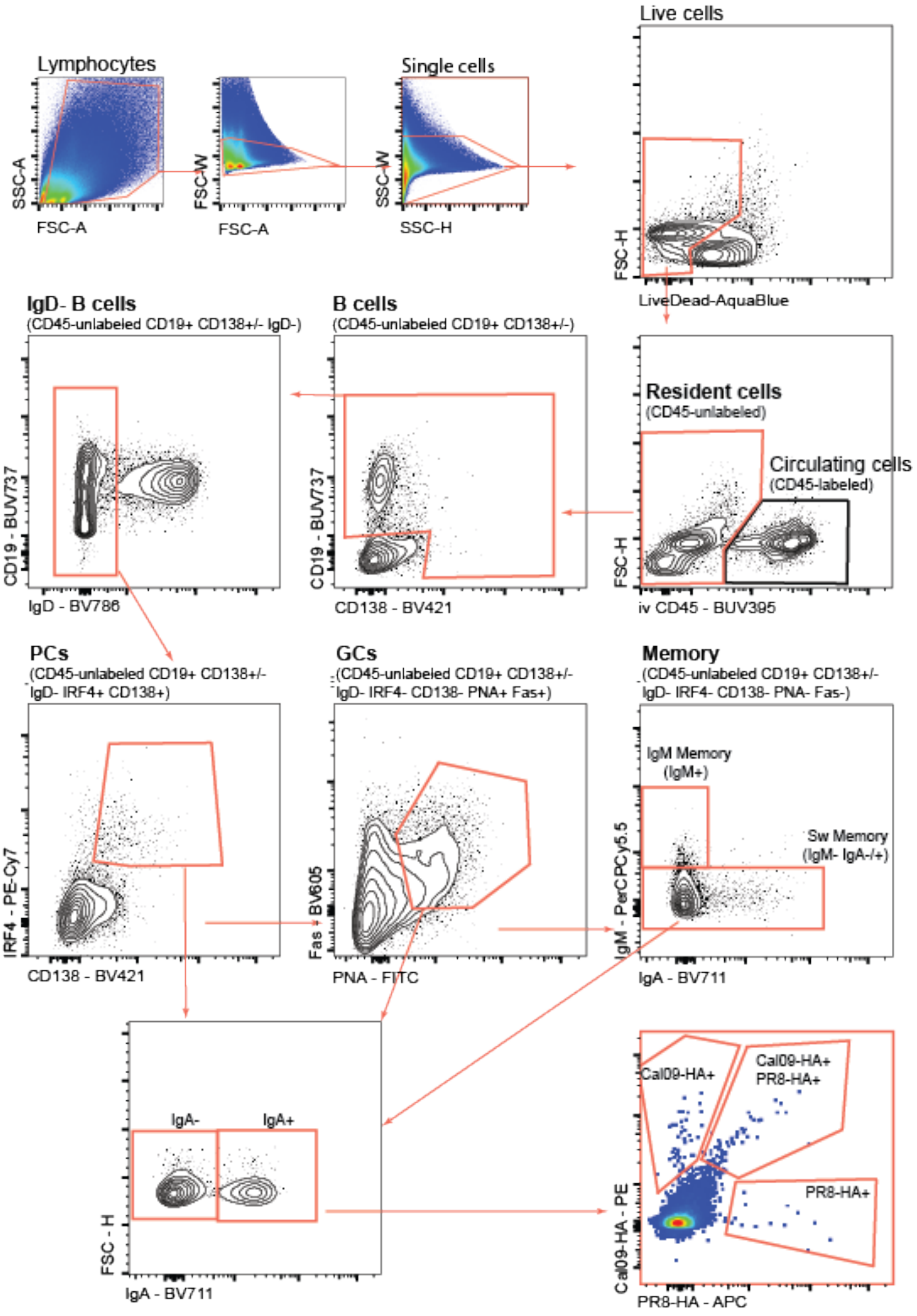


**Supplementary Figure 1.** *Gating strategy for lung and medLN B cells.*

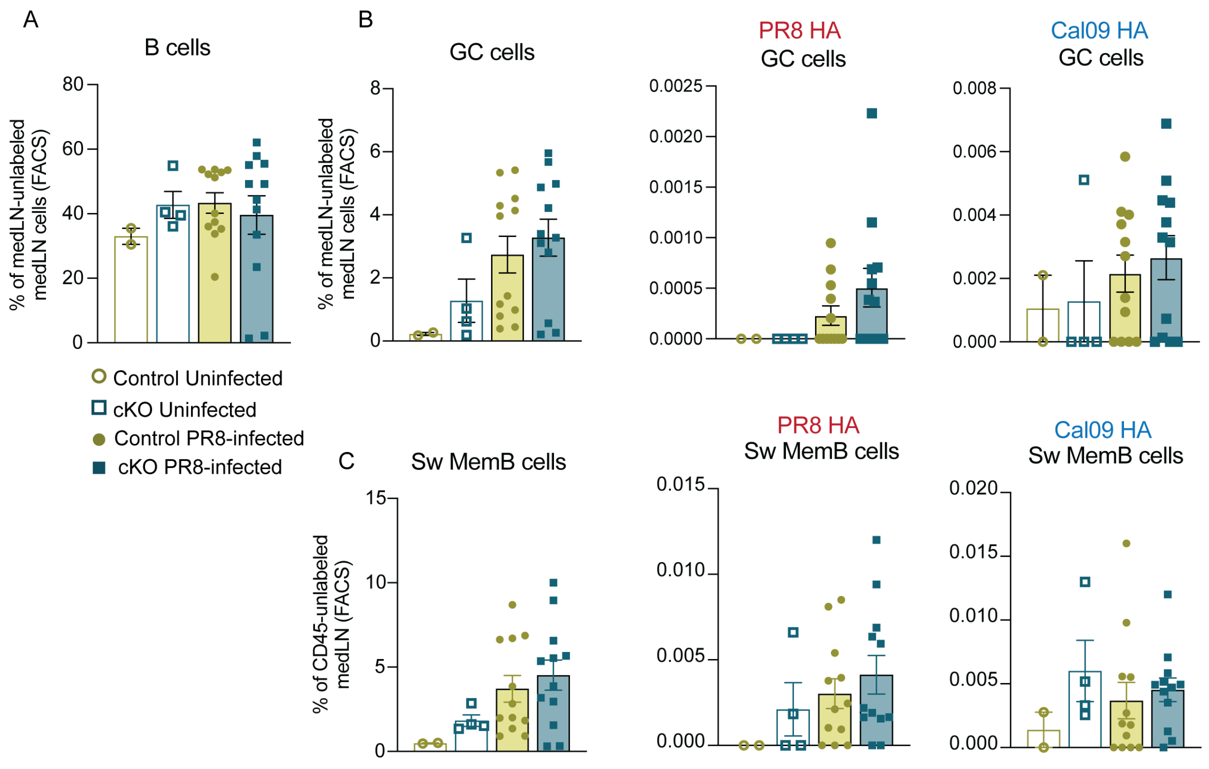


**Supplementary Figure 2.** $\alpha v^{+/+}$ $CD19^{Cre+}$ (control) and $\alpha v^{fl/fl}$ $CD19^{Cre+}$ (cKO) mice were infected i.n. with 100 plaque-forming units (PFU) of live H1N1 PR8 IAV. After 14 days of infection all mice received r.o. injection with 1μg α-CD45-BUV395 five minutes prior to euthanasia, lung and medLN were collected for analysis. (**A-B**) Quantification of the medLN B cells as a frequency of total resident medLN cells. **(B - C)** Quantification of the frequency of total GC **(B)** or switched memory **(C)** cells (left), PR8-HA specific (middle) and Cal09-HA specific (right) in the medLN. Each dot represents an individual mouse (n= 2mice for uninfected group and n=6-7 mice for infected groups). Data are means ± SEM from one representative experiment from 3 independent experiments. **p*<0.05 by Mann-Whitney U-test between the two PR8-infected groups.

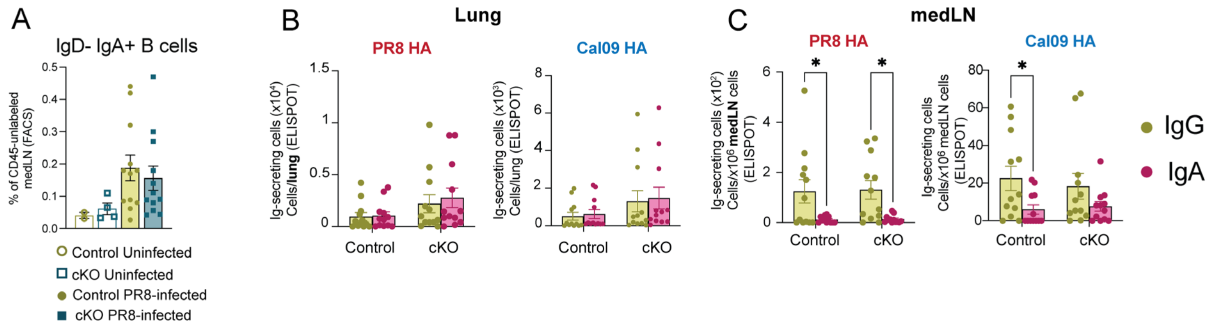


**Supplementary Figure 3.** $\alpha v^{+/+}$ $CD19^{Cre+}$ (control) and $\alpha v^{fl/fl}$ $CD19^{Cre+}$ (cKO) mice were infected i.n. with 100 plaque-forming units (PFU) of live H1N1 PR8 IAV. After 14 days of infection all mice received r.o. injection with 1µg α-CD45-BUV395 five minutes prior to euthanasia, lung and medLN were collected for analysis. **(A)** Quantification of medLN $IgD^-$ $IgA^+$ B cells by flow cytometry as a frequency of resident medLN cells. (**B-C**) Comparison of the Ig-secreting cells in the lungs **(B)** or medLN **(C)** that recognize PR8-HA (left) or Cal09-HA (right) as detected by ELISPOT. Each dot represents an individual mouse (n= 2mice for uninfected group and n=6-7 mice for infected groups). Data are means ± SEM of one representative experiment from 3 independent experiments. **p*<0.05 by Mann-Whitney U-test between the two PR8-infected groups.

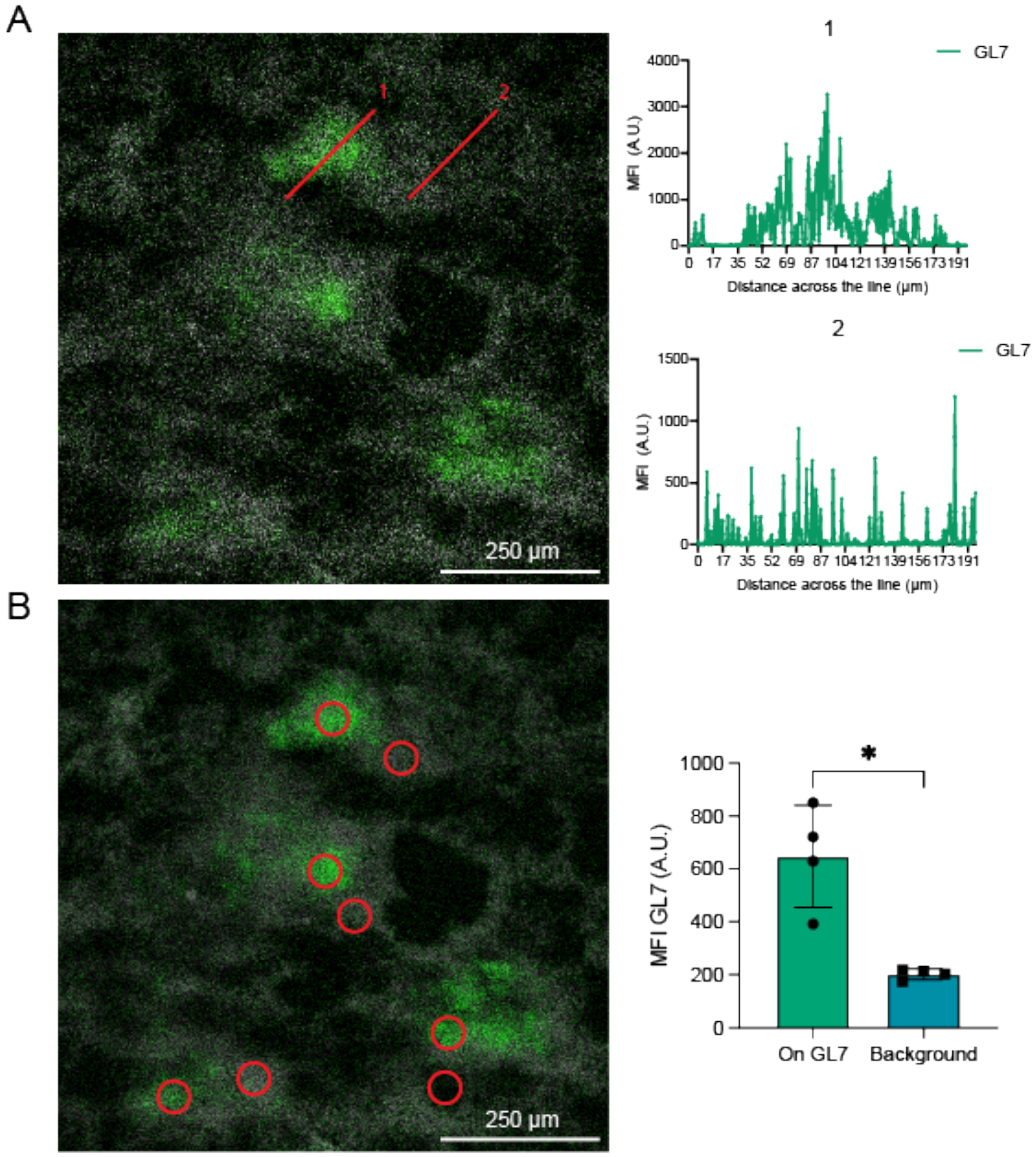


**Supplementary Figure 4.** *Identification of iBALT areas in the lung using GL7 as a marker for GC cells.* Mice were infected with live PR8 IAV as in **Figure 1A** and harvested for analysis of lungs by confocal microscopy as in **Figure 3A**. **(A)** Line scan across GL7 positive (1) and negative (2) region. Red lines represent areas delimited as GL7 staining or background. **(B)** Bar graph representing mean and SEM± of mean fluorescence intensity (MFI) of GL7 positive areas compared to background staining. Red circles represent areas delimited as GL7 positive or background. Each dot represent and individual are (GL7 stain or background) in one representative lung section. Scale bar 250 μm. *$p<0.05$ by Mann-Whitney U-test between the GL7 staining vs background.

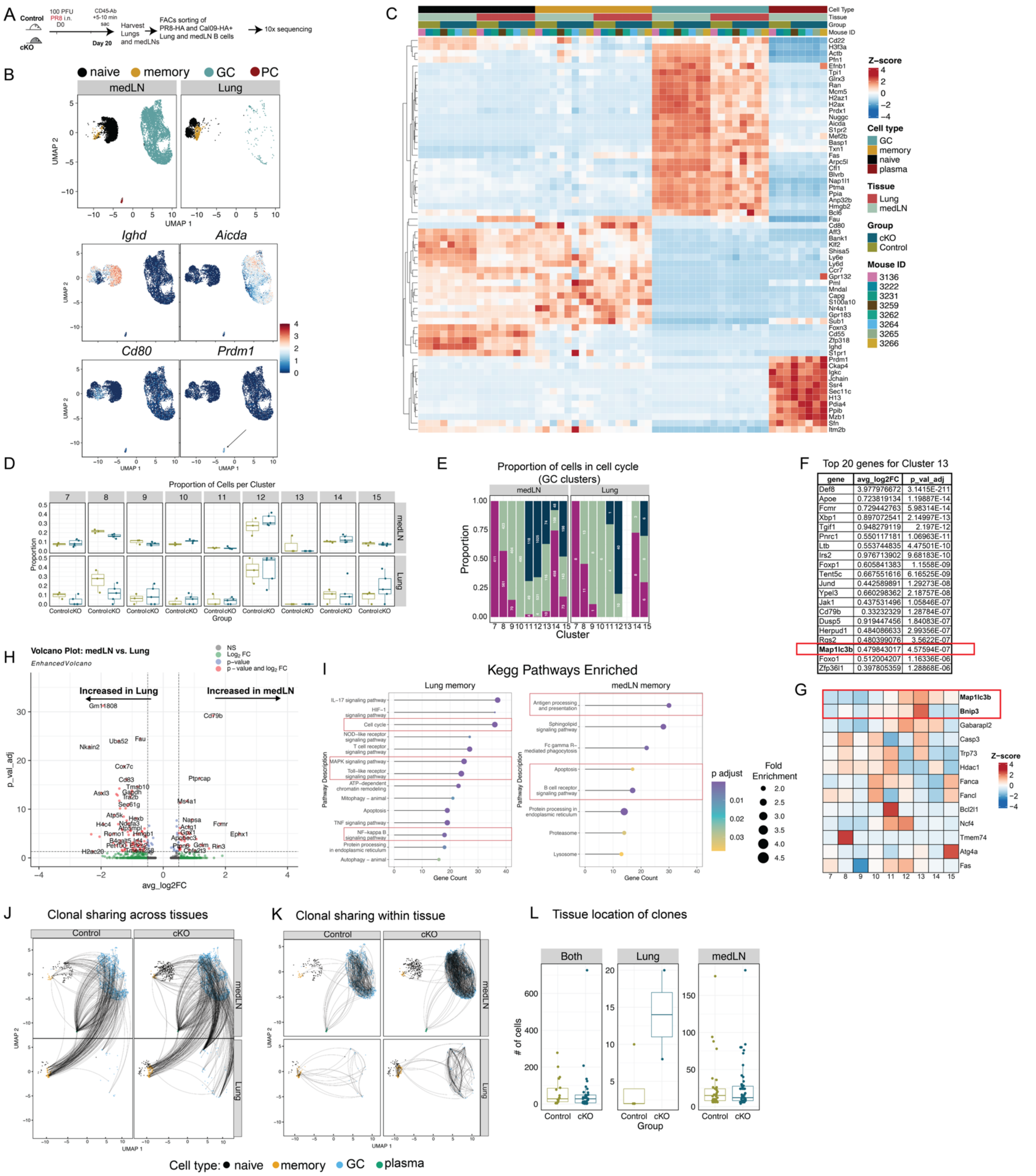


| gene | avg_log2FC | p_val_adj |
|---|---|---|
| Def8 | 3.977976672 | 3.1415E-211 |
| Apoe | 0.723819134 | 1.19887E-14 |
| Fcmr | 0.729442763 | 5.98314E-14 |
| Xbp1 | 0.897072541 | 2.14997E-13 |
| Tgif1 | 0.948279119 | 2.197E-12 |
| Pnrc1 | 0.550117181 | 1.06963E-11 |
| Ltb | 0.553744835 | 4.47501E-10 |
| Irs2 | 0.976713902 | 9.68183E-10 |
| Foxp1 | 0.605841383 | 1.1558E-09 |
| Tent5c | 0.667551616 | 6.16525E-09 |
| Jund | 0.442589891 | 1.29273E-08 |
| Ypel3 | 0.660298362 | 2.18757E-08 |
| Jak1 | 0.437531496 | 1.05846E-07 |
| Cd79b | 0.33232329 | 1.28784E-07 |
| Dusp5 | 0.919447456 | 1.84083E-07 |
| Herpud1 | 0.484086633 | 2.99356E-07 |
| Rgs2 | 0.480399076 | 3.5622E-07 |
| **Map1lc3b** | 0.479843017 | 4.57594E-07 |
| Foxo1 | 0.512004207 | 1.16336E-06 |
| Zfp36l1 | 0.397805359 | 1.28868E-06 |

**Supplementary Figure 5.** *Single cell RNA sequencing on antigen specific lung and medLN cells of mice infected with PR8 iAV.* $\alpha v^{+/+}$ $CD19^{Cre+}$ (control) and $\alpha v^{fl/fl}$ $CD19^{Cre+}$ (cKO) mice were infected with 100 PFU of live PR8 IAV. Lungs and medLN were harvested 20 days post infection. **(A)** Schematic of mice infection and tissue collection for lung and medLN sorting of antigen specific cells. **(B)** UMAP representation of scRNAseq data from B cells in lung and medLN. Clusters were assigned to one of four mayor B cell types based on the expression of the following markers: naïve (*Ighd*), GC (*Aicda*), Plasma cells (*Prdm1*), and memory (*Cd80*). **(C)** Heatmap of the z-score of canonical markers to differentiate various B cell subpopulations. **(D)** Box plot of the proportion of cells in each GC cluster in medLN (top) and lung (bottom) in the control and cKO mice. Each dot represents one mouse. **(E)** Quantification of the proportion of cells by cell cycle phase in the GC clusters. **(F)** Table of the top 20 genes with higher expression, as determined by adjusted p value. **(G)** Heatmap of the expression of GC autophagy-related genes. **(H - I)** Volcano plot comparing the genes highly expressed **(H)** and lollipop plot of the enriched KEGG pathways **(I)** in the lung memory (left) or the medLN memory (right). Lung memory was obtained by grouping clusters 0, 1 and 2-predominantely found in the lungs- and cluster 3 and 4-predominantely found in the medLN. **(J-K)** UMAP representation of the clonal sharing analysis comparing clones shared across tissues **(J)** and within same tissue **(K)** in the medLN and lungs of the control and cKO groups. Each dot represents a cell, each color represents a cell type, and the lines connect the clonal partners in different subpopulations and/or tissues. **(L)** Quantification of the number of GC cells per clone that are present in both tissues, only in the Lung or only in the medLN. Each dot represents a clone.

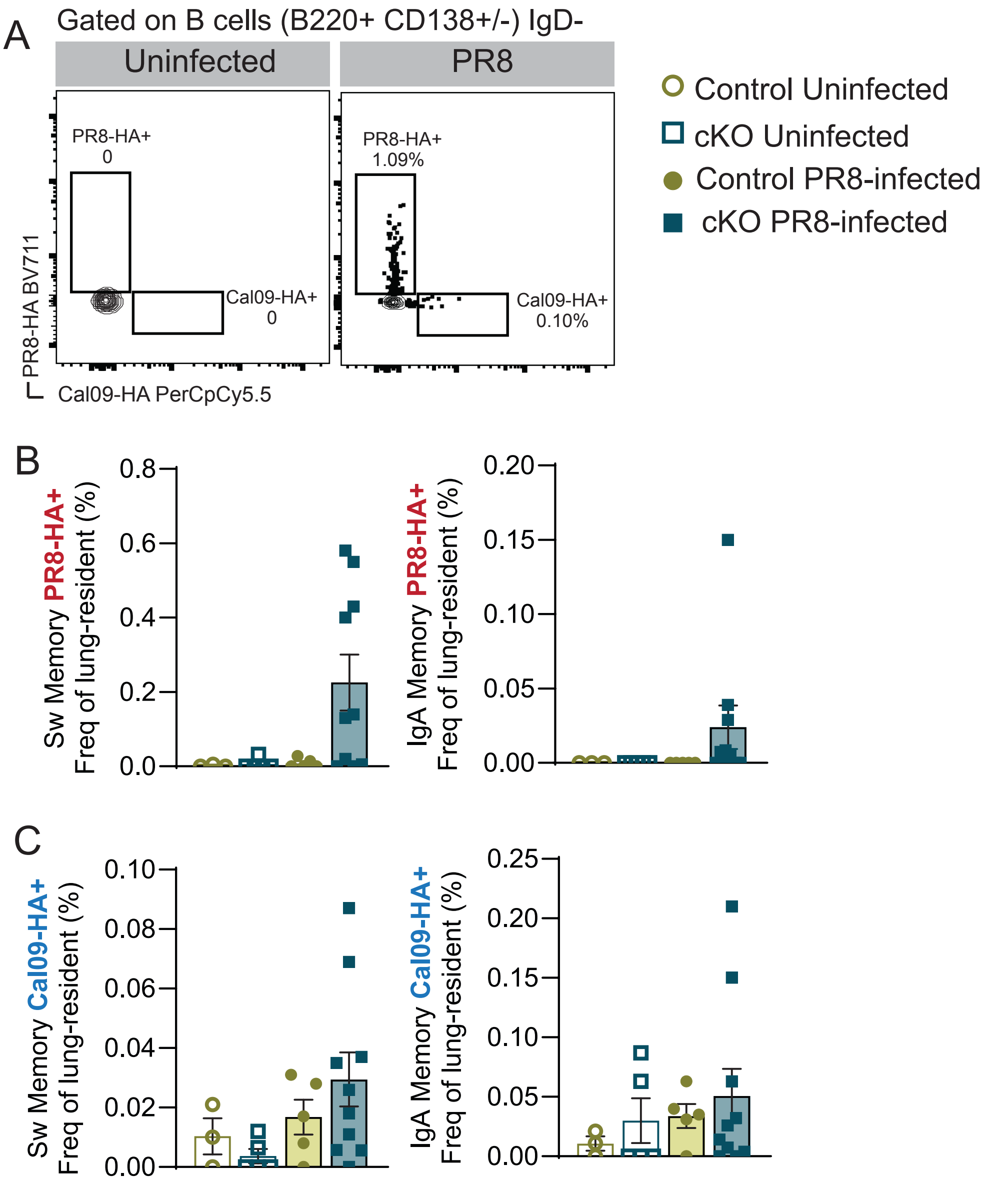


**Supplementary Figure 6.** *Absence of av in B cells leads to the differentiation of tissue resident cross reactive and antigen specific long lived memory B cells.* $\alpha v^{+/+}$ $CD19^{Cre+}$ (control) and $\alpha v^{fl/fl}$ $CD19^{Cre+}$ (cKO) mice were infected as in **Fig 6A**. **(A)** Representative flow cytometry gates of the PR8-HA – SA-BV711 and Cal09-HA – SA-PerCpCy5.5 tetramers in an uninfected (left) and infected (right) mouse; gated as live resident (CD45-unlabeled, see gating strategy on **Supp Fig1**) B220+ CD138+/- IgD-. **(B - C)** Quantification of the frequency of Sw and IgA memory specific for PR8-HA **(B)** or Cal09-HA **(C)** as identified by flow cytometry. Each dot represents one mouse; data are means SEM±. Representative experiment on 3 independent repeats.
